# Integration of evolutionarily distinct motility systems enables tunable cell propulsion

**DOI:** 10.1101/2025.10.29.685086

**Authors:** Anna Mas, Tâm Mignot, Marcelo Nollmann, Antoine Le Gall

## Abstract

Motility allows cells to explore their surroundings, evade threats, and coordinate collective behaviors. Many organisms, from bacteria to animals, encode more than one motility system, yet whether and how mechanistically distinct systems are integrated within individual cells remains largely unresolved. We address this question in the predatory bacterium *Myxococcus xanthus*, which navigates surfaces using two mechanistically and evolutionarily distinct motors: focal-adhesion-based gliding and type IV pili-driven twitching. For this, we develop an imaging-based quantitative behavioral phenotyping method to show that both motors are simultaneously active within individual cells and jointly power cell movement. Cells engaging both motors move faster and explore space more effectively than cells using either motor alone, expanding locomotion performance. Extracellular calcium tunes their relative contributions, progressively shifting cells from gliding-to twitching-dominated propulsion. Unexpectedly, the gliding motor can boost twitching even without anchoring to the surface, revealing that cooperation between motility systems extends beyond simultaneous force generation. Together, these findings reveal that evolutionarily distinct motility systems can function as components of a single, environmentally tunable propulsion strategy.

## INTRODUCTION

Cells adapt to changing environments by controlling when, where, and how they move, allowing them to explore space, escape threats, access nutrients, and coordinate with neighboring cells ^1–6^. While some cells rely on a single motility system, many organisms, including bacteria, archaea, protists, and higher eukaryotes, possess multiple, mechanistically and evolutionarily distinct propulsion mechanisms ^4^. For example, bacteria such as *Pseudomonas aeruginosa* can move by flagellar swimming or through type IV pili-dependent twitching motility ^7^. Similarly, the archaeon *Sulfolobus acidocaldarius* utilizes its archaella for swimming and pili for surface motility ^8^. Among eukaryotic microbes, *Salpingoeca rosetta* alternates between a flagellum-powered swimming form and an amoeboid crawling form ^9^. This coexistence of distinct motility systems is common and raises a key question: how are mechanistically distinct motility systems integrated within individual cells, and can their relative contributions be tuned to generate different behaviors? Answering this question requires approaches capable of resolving the functional contribution of each system simultaneously and non-invasively within individual living cells, a technical challenge that has so far limited progress in this area.

The soil bacterium *Myxococcus xanthus* provides a powerful model to address this question. *M. xanthus* is a predatory bacterium that must navigate complex surface environments to locate and consume prey, often coordinating its movements across large multicellular groups ^10–13^, using two mechanistically distinct motility systems ^10,14^. A-motility (adventurous gliding) is powered by a proton-motive-force motor (AglRQS) and organized by the cytosolic scaffold AglZ ^15^, engaging the CglB adhesin to form focal-adhesion-like sites that anchor the cell to the substrate and allow the motor to pull the cell forward ^16,17^. In contrast, S-motility (social twitching) relies on type IV pili: PilA subunits are polymerized into extracellular filaments by the PilB ATPase at the leading pole, attach to surfaces, and retract via the PilT ATPase to drag the cell forward ^18,19^. These two systems are evolutionarily and mechanistically unrelated, and their downstream activation remains distinct ^20,21^, yet they share a single polarity control module: MglA-GTP recruits both complexes to one pole, while the Frz chemosensory pathway relocalizes them to the opposite pole to trigger reversals ^22,23^.

Early genetic work by Hodgkin and Kaiser established that each system is encoded by a largely distinct gene group, with A-motility genes associated with single-cell gliding and S-motility genes associated with collective cell movement, providing separable genetic and phenotypic readouts that have anchored the study of A-and S-motility for decades ^24,25^. In contrast, recent work showed that single-system mutants are not confined to adventurous and collective cell movements, and that wild-type cells dynamically transition between these cell groups ^26^. These results are inconsistent with the exclusive association of A-and S-motors with adventurous and social motility, and instead suggest that their activities may be coordinated within individual cells rather than segregated between distinct cellular states.

To determine whether and how these two motility systems are integrated within individual cells, we combined high-resolution single-cell tracking, behavioral phenotyping, and agent-based modeling. Remarkably, we demonstrate that A-motility drives both single and collective cell movement, inconsistent with twitching being exclusively responsible for social behavior. Next, we show that both motility systems are simultaneously deployed in single cells. By applying behavioral phenotyping, we conclude that both motors can actively propel single cells, and that extracellular calcium progressively shifts cells from gliding-to twitching-dominated propulsion, thereby tuning the relative contributions of the two systems. This combined action of A-and S-motility motors allows cells to move faster and explore space more effectively than single-motility mutants, an enhancement that cannot be explained by the additive contribution of each system alone. Finally, we show that the gliding motor can enhance twitching motility independently of substrate anchoring, indicating that cooperation between the two systems extends beyond simultaneous force generation. Together, these findings show that single cells can integrate evolutionarily distinct motility systems to generate environmentally tunable propulsion.

## RESULTS

### A-Motility Is Propulsive Across All Cell Contexts During Predation

To determine whether the two motility systems can be integrated within individual cells, we first asked whether they are deployed in the same cellular contexts. Previous models proposed that A-motility and S-motility are segregated at the population level, with A-motility restricted to solitary ’scout’ cells at the colony edge and S-motility dominating collective swarms. If this segregation holds, co-engagement within the same cell would be structurally impossible. We therefore considered three possible modes of motility system deployment: cells where only one system is present (mutually exclusive deployment), cells where both are present but only one is active at a time (co-deployment with alternating propulsion), or cells where both are present and simultaneously active (co-deployment with co-propulsion) (Fig. 1a). To distinguish between these models, we combined complementary imaging strategies (Fig. 1b, Fig. S1a). Phase-contrast imaging was used to detect and track the individual cells positions at high spatiotemporal resolution (Fig. 1b, Fig.S1b), while a combined TIRF/HILO fluorescence imaging system allowed us to monitor multiple fluorescent reporters simultaneously to detect the presence and activity of A and S-motility systems in single cells (Fig. 1b; Fig. S1a, Movie S1).

**Figure 1:**
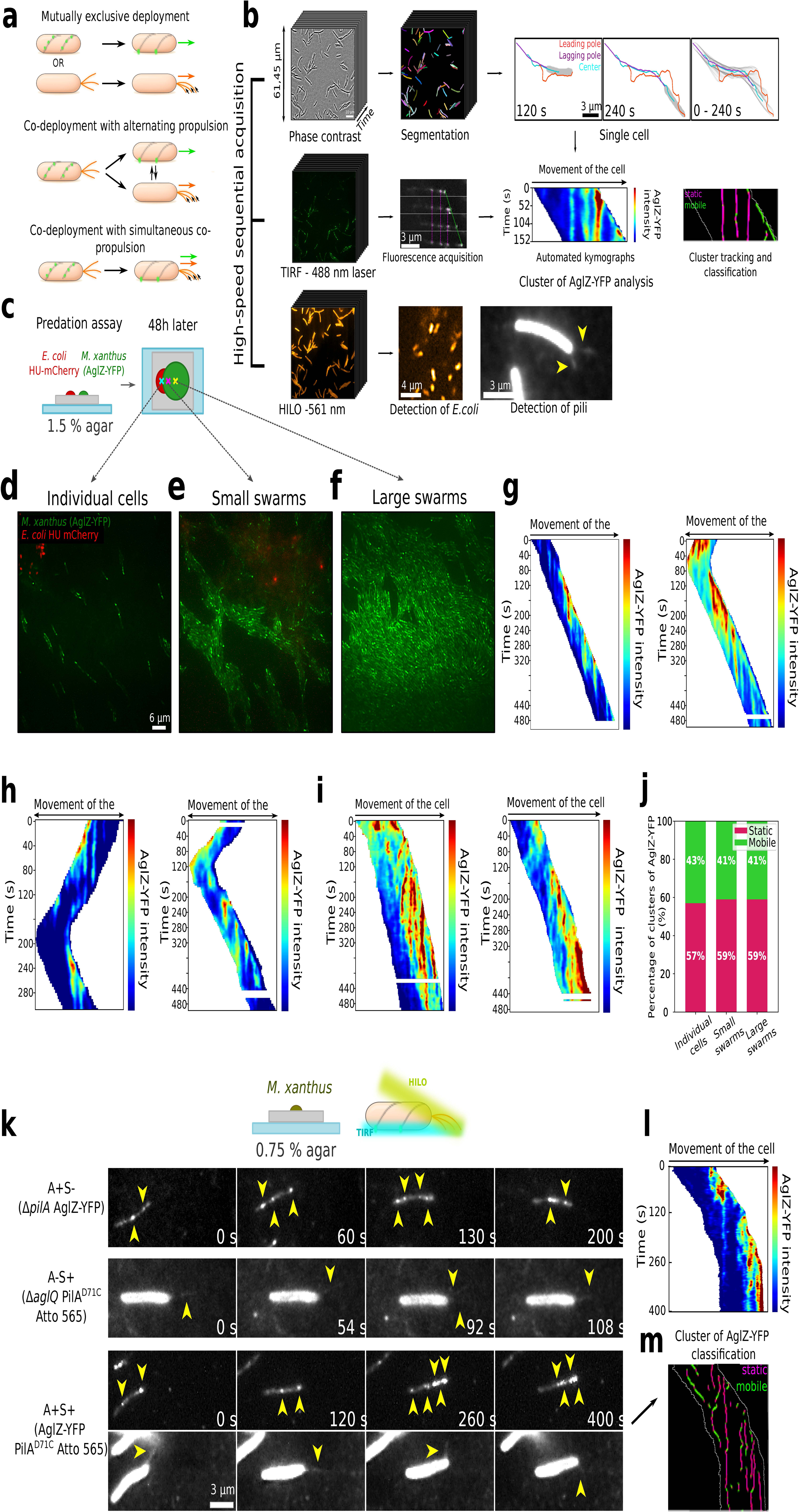
A-Motility Is Propulsive in Isolated and Swarming Cells and Co-Deployed with S-Motility in Individual Cells. **(a)** Models for the deployment and usage of motility systems in *Myxococcus xanthus* cells. Top: Exclusive deployment model, cells deploy and propel using only one motility system at a time. Middle: Co-deployment with switchable propulsion model, both systems are deployed within the same cell, but only one is propulsive at a time. Bottom: Co-deployment and co-propulsion model, both systems are deployed and simultaneously contribute to propulsion within the same cell. Green: Agl-Glt complexes (transient on the left; fixed on the substrate on the right), Orange: Type IV pili. **(b)** Top row: Phase-contrast imaging & single-cell tracking. A rapid time series (left) is segmented to identify individual cells (middle), whose leading pole (red), lagging pole (purple), and center (cyan) are tracked over time (right). Representative trajectory overlays are shown for 120 s and 240 s, as well as the full 0-240 s interval. Middle row: TIRF imaging of AglZ-YFP clusters & kymograph analysis. Cells expressing AglZ-YFP are imaged by TIRF (488 nm) (left) and cluster fluorescence along each cell body is extracted into automated kymographs (middle), where the x-axis corresponds to distance along the cell and the y-axis to time. Clusters are then classified as static (magenta) or mobile (green) by tracking their intensity peaks over successive frames (right). Bottom row: HILO detection of prey and pili (left). A simultaneous HILO time series (561 nm) detects fluorescently labeled *E. coli* prey (middle) in predation assays or type IV pili (yellow arrowheads) in motility assays emanating from *M. xanthus* cells (right). **(c)** Schematic of the predation assay where A⁺S⁺ *M. xanthus* cells expressing AglZ-YFP (green) are spotted next to *E. coli* HU-mCherry prey on 1.5% agar. After 48 h of co-incubation, distinct populations, large swarms, small swarms, and individual cells, were identified and imaged for downstream cluster analysis. **(d-f)** Fluorescence images of A⁺S⁺ AglZ-YFP (green clusters) and *E. coli* HU-mCherry (red) in the three depicted zones of panel (c) of the predation assay: isolated cells (d), small swarms (e), and large swarms (f). **(g-i)** Automated kymographs of AglZ-YFP signal in A⁺S⁺ cells extracted from panels (d-f). Two example cells are shown for each condition (d: individual cells, e: small swarms, f: large swarms). **(j)** Bar chart showing the percentage of AglZ-YFP clusters classified as static (magenta) or mobile (green) across the entire field of view in each population type. **(k)** Time-lapse HILO/TIRF of pili and AglZ-YFP dynamics in different genetic backgrounds on 0.75% agar pads. Top: A⁺S⁻ (Δ*pilA* AglZ-YFP) Middle: A⁻S⁺ (Δ*aglQ* PilA^D71C^ Atto 565) Bottom: A⁺S⁺ (AglZ-YFP, PilA^D71C^ Atto 565). Yellow arrowheads indicate AglZ-YFP clusters (top), fluorescent pili (middle), or both (bottom). **(l)** Automated kymographs of the A⁺S⁺ cell shown in panel (k). **(m)** Cluster classification of static (magenta) or mobile (green) clusters of AglZ-YFP from the same A⁺S⁺ cell, overlaid on the kymograph trace (white outline).

To directly monitor A-motility activity, we tagged AglZ, a cytoplasmic component of the Agl-Glt machinery, with YFP and visualized it in TIRF mode to reveal substrate-anchored focal adhesions; static AglZ-YFP clusters mark propulsive adhesion sites ^16,17^. In a predation assay (Fig. 1c), wild-type cells expressing AglZ-YFP were plated with prey on nutrient-poor agar and imaged after 48h of co-incubation, a condition that generates spatially heterogeneous cell contexts, including both isolated and collective behaviors, and thus provides a direct test of whether A-motility is restricted to specific subpopulations ^26^. As expected, AglZ-YFP clusters were detected in isolated cells at the colony front (Fig. 1d). Strikingly, such clusters were also present in small swarms (Fig. 1e) and in large swarms behind the front (Fig. 1f). To assess whether these clusters were propulsive, we examined multiple cells from each population type: kymographs of AglZ-YFP intensity consistently exhibited vertical streaks (Fig. 1g-i), the hallmark of static, substrate-anchored adhesions. Such vertical streaks arise because AglZ-containing focal adhesion complexes remain fixed relative to the substrate while the cell body moves forward, a defining feature of propulsive A-motility ^17^. Clusters were classified as ‘static’ or ‘mobile’ using an unsupervised Gaussian mixture model trained on multiple motility features, including instantaneous velocity and trajectory geometry (see Supplementary Methods). Quantitative tracking confirmed that over 41% remained immobile relative to the substrate (Fig. 1j). Together, these data demonstrate that A-motility is actively deployed and propulsive in both isolated cells and high-density swarms where S-motility was thought to dominate. While we expect that S-motility will contribute to, or even drive, collective cell motion within high-density swarms, high cell densities prevented direct visualization of S-motility structures within these dense structures. To directly assess whether both motility systems can be deployed within the same individual cells, we therefore turned to a more controlled single-cell imaging setup.

### Both Motility Systems Are Co-Deployed in Individual Cells

To directly monitor motility system deployment at the single-cell level, we performed high-resolution imaging of isolated cells deposited on a 0.75 % agar pad (an intermediate concentration chosen to promote both systems), covered with a coverslip. To directly detect if both motility systems are present in single cells, we imaged wild-type cells co-expressing AglZ-YFP and fluorescently labeled pili. For the latter, PilA, the major type IV pilin, was labeled with Atto 565 and imaged under HILO illumination to detect pili within ∼1 μm of the surface ^21^. Remarkably, in wild-type cells, focal adhesions and Atto 565-labeled pili were often detected in the same single cells (Fig. 1k; Movies S3-S4), directly demonstrating that A-and S-motility systems are co-deployed. Kymograph and cluster tracking of AglZ-YFP (Fig. 1l-m, Fig. S1c) confirmed that adhesion sites in these wild-type cells were propulsive. To verify that each motility system can function on its own, we also imaged motility mutants where either A-motility (A⁻S⁺) or S-motility (A⁺S⁻) were inactive. In the A⁺S⁻ mutant (Δ*pilA*), AglZ-YFP clusters were clearly visible and associated with cell propulsion, confirming that A-motility remained fully deployed and functional under these conditions (Fig. 1k; Fig. S1d). In contrast, the A⁻S⁺ mutant (Δ*aglQ*) deployed pili at its leading pole but did not move, indicating that S-motility machinery can be deployed without necessarily driving propulsion in isolated cells (Movie S5). Notably, wild-type cells (A⁺S⁺) moved slightly faster than A⁺S⁻ mutants, with no obvious differences in the relative movements of the leading and lagging poles, a proxy for “cell pole behaviors” that will be analyzed in detail below (Fig. S1e-g). This observation raised the possibility that pili may augment propulsion when A-motility is active, though with only a small effect on overall cell movement on agar. In summary, these experiments establish that both motility systems can be deployed within the same single cell. However, whether they also operate simultaneously to drive propulsion remains unresolved, as S-motile cells are poorly motile under these conditions. Moreover, the mere presence of pili does not demonstrate active propulsion, as pili can serve other functions including DNA uptake and surface sensing ^27^. Directly resolving their propulsive contribution therefore requires both a substrate supporting processive motility by both systems and a quantitative approach capable of distinguishing active twitching from passive pilus deployment.

### A-and S-motility simultaneously propel individual cells

To directly test if both systems can simultaneously contribute to propulsion within single cells, we implemented a setup where both A⁺S⁻ and A⁻S⁺ mutants are able to move processively. Specifically, we employed a microfluidic chamber with coverslips coated with chitosan, a polysaccharide that promotes A-motility ^28^. Under these conditions, both motility systems remain co-deployed (Movie S6) as in agar, and both A⁺S⁻ and A⁻S⁺ mutants were able to move (Fig. S2a), confirming that chitosan-coated surfaces support processive motility by either system. This provided the conditions needed to ask whether A⁺S⁺ cells can engage both motility systems simultaneously within the same cell.

Notably, joint detection of AglZ-YFP foci and of pili in A⁺S⁺ cells indicates that both motilities are simultaneously deployed on chitosan-coated coverslips (Fig. 2a, Movie S7). To determine if A-motility remains propulsive on this substrate, we analyzed AglZ-YFP clusters using kymographs (Fig. 2b) and cluster tracking (Fig. 2c). These analyses showed that many AglZ-YFP foci remained stationary relative to the substrate, consistent with active focal adhesions throughout the A⁺S⁺ population. Fluorescently-labeled pili localized at the leading pole of the same cells displaying static AglZ-YFP clusters, demonstrating that both A and S motilities can be co-deployed. Interestingly, quantification of the number of static AglZ-YFP clusters per cell was similar in A⁺S⁺ and A⁺S⁻ strains (Fig. 2d; Fig. S2b), indicating that pili do not affect the assembly of focal adhesion complexes under these conditions.

**Figure 2.**
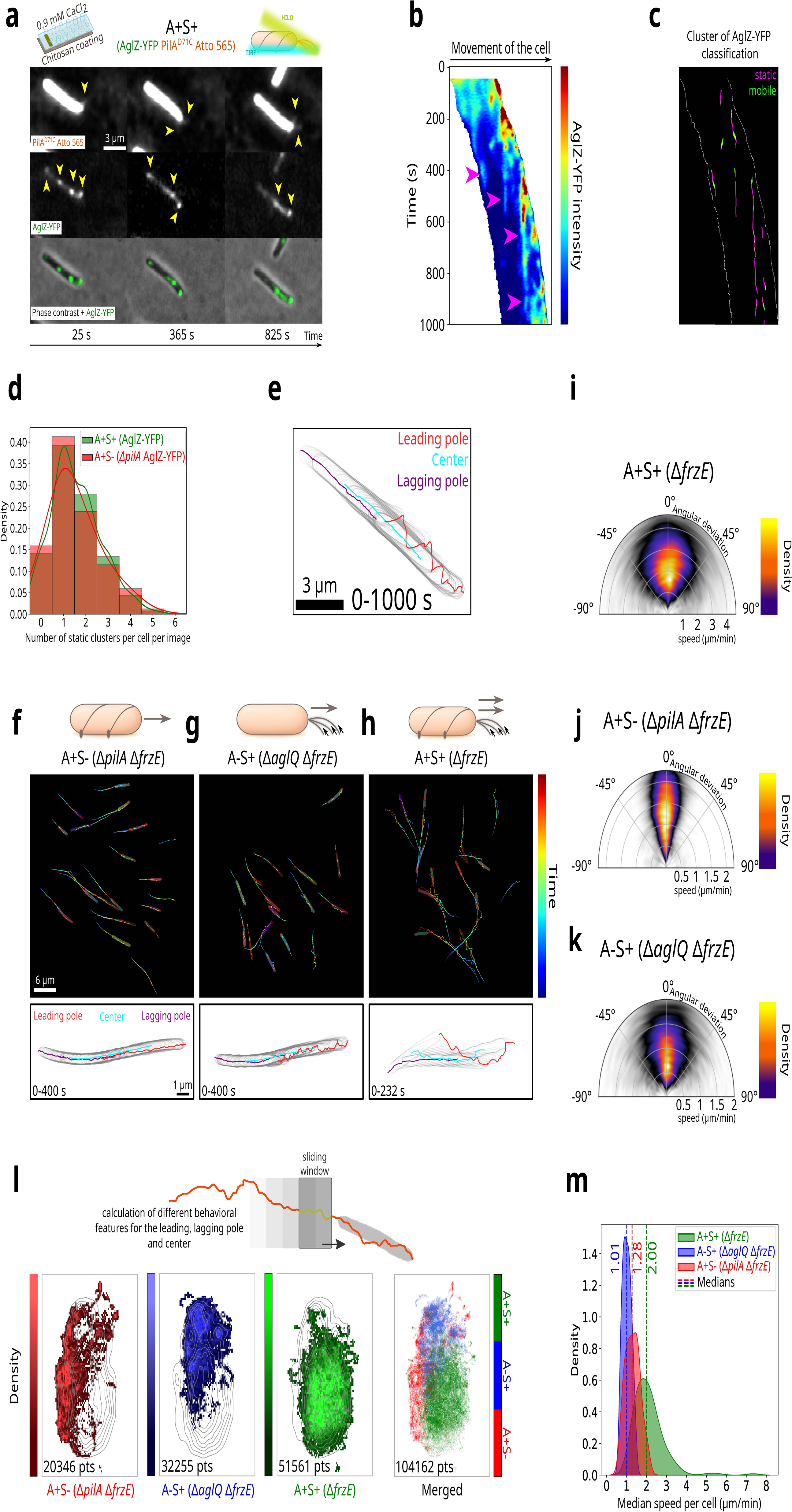
: Co-Propulsion by A-and S-Motility Generates Distinct Behaviors and faster movement within Single Cells. **(a)** Combined TIRF (488 nm) and HILO (561 nm) fluorescence images of A⁺S⁺ *M. xanthus* coexpressing AglZ-YFP and PilA^D71C^ Atto 565 on a chitosan surface with 0.9 mM CaCl₂. Top row: HILO channel showing labeled pili (yellow arrowheads). Middle row: TIRF channel showing AglZ-YFP clusters. Bottom row: merged phase-contrast and AglZ-YFP for cell outline. **(b)** Automated kymograph of AglZ-YFP intensity extracted along the long axis of the same A⁺S⁺ cell. Magenta arrowheads mark static clusters. **(c)** Cluster classification for the cell in (A-B): static AglZ-YFP clusters (magenta) versus mobile clusters (green) over the 0-1000 s interval, overlaid on the kymograph trace (white outline). **(d)** Distribution of static AglZ-YFP clusters per cell per image for A⁺S⁻ (Δ*pilA*, red, N = 2 replicates, n = 28 cells) and A⁺S⁺ (green, N = 2 replicates, n = 123 cells) strains. Bars show counts; solid lines are kernel density estimates. **(e)** Trajectories of the leading pole (red), lagging pole (purple), and cell center (cyan) for the A⁺S⁺ cell from panels (a-c) over 0-1000 s. Gray region indicates the cell contour projected through time. **(f-h)** Full-field trajectories (top, over 597 s) colorcoded by time and single-cell examples (bottom, over 400 s) showing leading pole (red), lagging pole (purple), and center (cyan) paths for: (f) A⁺S⁻ (Δ*pilA* Δ*frzE*), (g) A⁻S⁺ (Δ*aglQ* Δ*frzE*), (h) A⁺S⁺ (Δ*frzE*). Cell outlines (light gray) at each timepoint highlight contour evolution. **(i)** Polar density plot of instantaneous cell speed (radial axis, µm/min) versus angular deviation of the leading pole (0° = forward; ±90° = lateral) for A⁺S⁺ (Δ*frzE*; N=4, n=159). Color scale indicates point density. **(j-k)** Polar density plots as in (i) for: (j) A⁺S⁻ (Δ*pilA* Δ*frzE*; N=3, n=61), (k) A⁻S⁺ (Δ*aglQ* Δ*frzE*; N=3, n=67). **(l)** UMAP projections of 48 spatial behavior features (excluding any speed metrics), computed for the leading pole, center, and lagging pole over sliding time windows, shown as a two-dimensional density map. Above each density map is a schematic of a single-cell leading-pole trajectory (orange) with a shaded time window, illustrating that each “behavior segment” corresponds to one such window along the leading-pole path, from which 16 spatial features are extracted (48 total when including the center and lagging pole, not shown). Each small square (bin) in the map represents a region of UMAP space, and its color indicates how many “behavior segments” fall into that bin (lighter = more segments). Projections are shown separately for A⁺S⁻ (Δ*pilA* Δ*frzE*, red), A⁻S⁺ (Δ*aglQ* Δ*frzE*, blue), and A⁺S⁺ (Δ*frzE*, green), with the far-right panel overlaying all three densities. Iso-contours in the first three panels indicate the combined occupancy across strains. **(m)** Kernel density distribution of the median instantaneous speed per cell for A⁺S⁺ (Δ*frzE*, N = 4 replicates, n = 167 cells), A⁺S⁻ (Δ*pilA* Δ*frzE*, N = 3, n = 61 cells), and A⁻S⁺ (Δ*aglQ* Δ*frzE*, N = 3, n = 67 cells) strains. Dashed line indicates population medians.

To determine whether pili also contribute to propulsion in A⁺S⁺ cells, one would need to correlate individual pilus extension-retraction events with cell movement. In practice, however, pilus fluorescence is dim and photobleaches rapidly, and its fast kinetics of retraction demand both high frame rates and light doses that lead to phototoxicity. These constraints make it impractical to directly link pilus retraction events to cell propulsion, so we instead sought a robust signature of active pilus retraction. By first examining the trajectories of the leading pole, the cell center, and the lagging pole of A⁺S⁺ cells, we noticed pronounced deflections of the leading pole relative to the cell axis (Fig. 2e; Fig. S2c, Movie S8), suggesting that pili might pull the leading pole off-axis even in the presence of stationary focal adhesions. To minimize the confounding effect of cellular reversals, we tracked the same three trajectory positions in Δ*frzE* backgrounds, in which deletion of the FrzE chemosensory kinase abolishes directional reversals. In A⁺S⁻ (Δ*pilA*, Δ*frzE*) cells, both the leading and lagging poles remained tightly aligned with the cell center, yielding straight trajectories (Fig. 2f, Movie S9). By contrast, A⁻S⁺ (Δ*aglQ*, Δ*frzE*) cells displayed pronounced leading-pole deflections while their lagging poles stayed aligned, confirming that pilus activity alone can shift the leading pole off-axis (Fig. 2g, Movie S9). Remarkably, A⁺S⁺ (Δ*frzE*) cells also displayed large leading-pole fluctuations while maintaining lagging-pole alignment (Fig. 2h, Movie S9).

To quantify these deflections, we measured the angle distributions between the leading pole and the cell axis. In A⁺S⁺ cells, these angles reached up to ∼45° (Fig. 2i, Fig. S2d). In contrast, the leading poles of A⁺S⁻ cells fluctuated in a considerably narrower range (∼20°, Figs. 2j, S2d), consistent with the activity of pili being responsible for these deflections. As expected, the leading pole of A⁻S⁺ cells deflected with similar amplitudes to A⁺S⁺ cells (∼40°, Fig. 2k, Fig. S2d), in agreement with large off-axis deviations of the pole being driven by pilus activity. Altogether, these data show coexisting stationary AglZ-YFP foci and pronounced pole fluctuations, therefore supporting the view that focal adhesions and pili are not only co-deployed but also jointly power single-cell propulsion across the A⁺S⁺ population.

Next, we investigated how co-deployment of both systems impacts single-cell movement by extracting 16 local motility descriptors, such as tortuosity, local radius of gyration, directional autocorrelation, and angular dispersion, along each of the leading-pole, cell-center, and lagging-pole trajectories (48 features in total, see Supplementary Material). To remove any time dependence, all trajectories were first rediscretized at uniform distance intervals. These descriptors therefore captured how each tracked position of the cell samples its local environment in terms of path geometry, directionality, and area explored. Importantly, none of the 16 features included time-derivative information such as velocity or acceleration, ensuring that these features focused strictly on how cells explore space rather than how fast they move.

To visualize and compare these complex, high-dimensional behavioral profiles, we projected the features into two dimensions using Uniform Manifold Approximation and Projection (UMAP), a nonlinear dimensionality reduction method that preserves local structure ^29^. In this projected “behavioral space,” each point corresponds to a segment of a cell trajectory described by its local motility features, so that proximity in the plot reflects similarity in movement patterns. Data points from A⁺S⁻, A⁻S⁺, and A⁺S⁺ cells formed largely distinct clusters (Fig. 2l; Fig. S2e-h), with only minimal overlap between strains, indicating that each strain exhibits a characteristic “behavioral fingerprint” defined by its local motility patterns. Notably, this segregation emerged without providing any information about the cell genotype, indicating that strain-specific behaviors can be reliably distinguished based solely on local spatial features of cell motion. A⁺S⁺ cells occupied a clearly separate region of the behavioral space, distinct from both A⁺S⁻ and A⁻S⁺ mutants, confirming that they exhibit a unique motility signature that differs from either single-motility system alone.

Finally, we compared the speeds of the three strains to evaluate the functional impact of co-existing motility systems. Remarkably, A⁺S⁺ cells moved approximately twice as fast as either single-motility mutant (Fig. 2m; Fig. S2i), indicating that co-propulsion results not only in behavioral diversification but also in enhanced locomotion capacity. Thus, integration of the two systems generates behavioral and locomotor outputs distinct from those produced by either system alone. This raised the question of why A⁺S⁺ cells move faster. Does co-propulsion simply sum forces from both systems, or is their activity regulated?

### Extracellular Calcium Tunes a Continuum of Motility Regimes

To test whether the relative contributions of A-and S-motility can be tuned, we varied extracellular CaCl₂ which is known to modulate pili activity via PilY1-dependent assembly and stabilization ^30,31^, while noting that calcium may also affect A-motility ^32^.

We leveraged the ability of our microfluidic system to deliver buffers to the same cell cohort to progressively expose the same cells to increasing CaCl₂ concentrations (from 0 to 4 mM) while measuring the impact to cell velocities and behavior. In the A⁺S⁻ mutant, cells remained motile at all CaCl₂ levels, with the leading/lagging poles and the cell center tracing straight, tightly aligned paths (Fig. 3a; Movie S10). At 0 mM CaCl₂, cells displayed short trajectories, which lengthened at moderate calcium concentrations before shortening again at higher levels, suggesting a non-linear effect of calcium on A-motility. By contrast, the A⁻S⁺ mutant was immotile below ∼0.7 mM CaCl₂ and then displayed a sharp activation of motility, exhibiting processive movement and growing pole fluctuations at higher CaCl₂ concentrations (Fig. 3b, Movie S11). Overall, these results indicate that calcium activates S-motility in line with previous studies ^31^, but also confirm that CaCl₂ modulates A-motility behavior ^32^.

**Figure 3.**
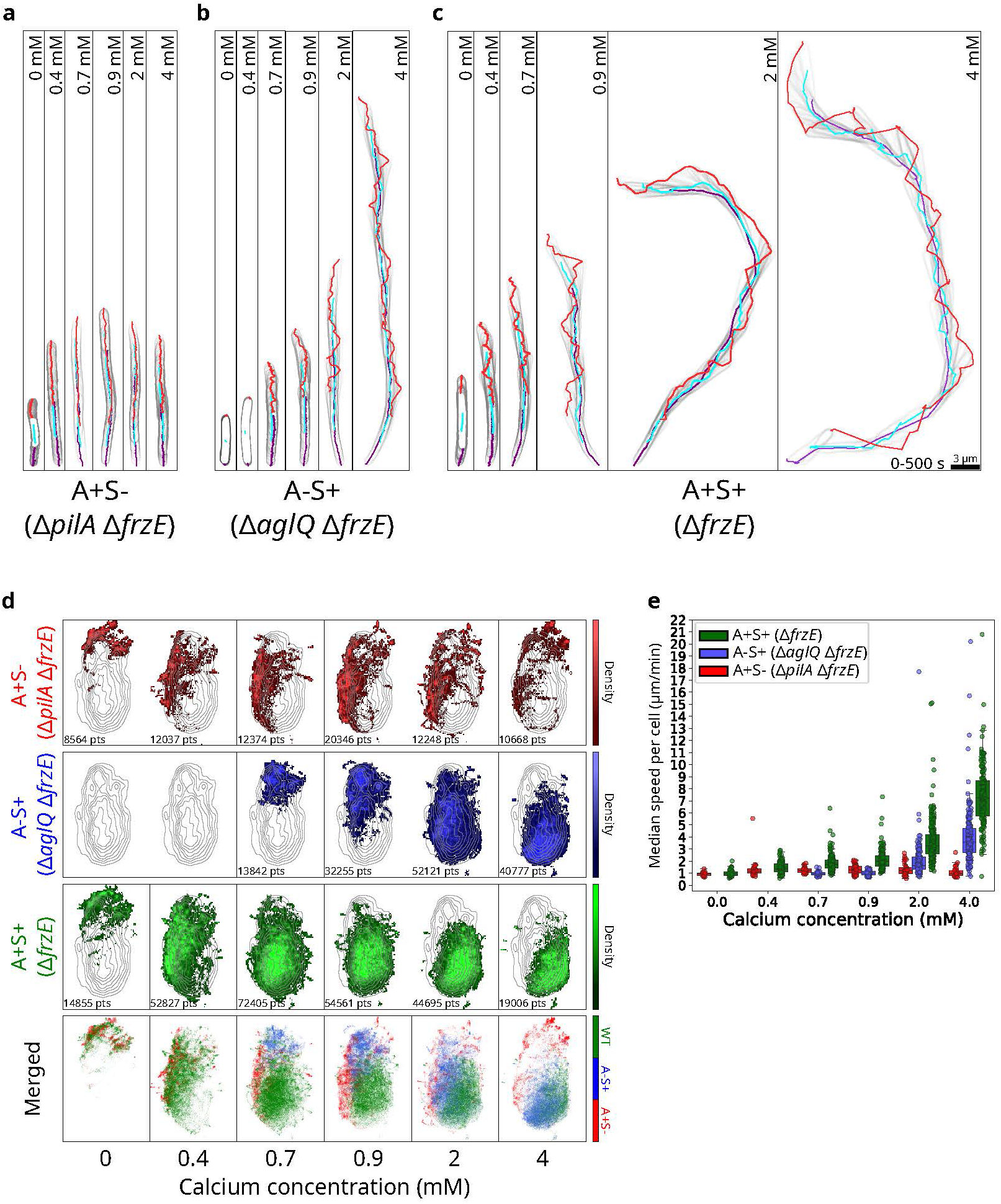
: Motility Behavior and Propulsion Dynamics Are Modulated by Environmental Conditions. **(a-c)** Representative single-cell trajectories of the leading pole (red), lagging pole (purple), and cell center (cyan) under increasing CaCl₂ concentrations (0, 0.4, 0.7, 0.9, 2, 4 mM) for: (a) A⁺S⁻ (Δ*pilA* Δ*frzE*), (b) A⁻S⁺ (Δ*aglQ* Δ*frzE*), (c) A⁺S⁺ (Δ*frzE*). Light-gray outlines show cell contours at each timepoint (light gray). **(d)** UMAP embeddings of the same 48 spatial behavior features introduced in Figure 2l, here shown as density maps for each strain across the CaCl₂ series. From top to bottom: A⁺S⁻ (Δ*pilA* Δ*frzE*, red), A⁻S⁺ (Δ*aglQ* Δ*frzE*, blue), and A⁺S⁺ (Δ*frzE*, green), and merged. The shifting density clouds illustrate how each strain’s repertoire of behaviors moves through feature space as calcium increases. Iso-contours represent the combined behavioral density of all three strains pooled at the calcium concentration of 0.9 mM. **(e)** Median instantaneous speeds per cell across CaCl₂ concentrations for A⁺S⁺ (Δ*frzE*, green), A⁺S⁻ (Δ*pilA* Δ*frzE*, red), and A⁻S⁺ (Δ*aglQ* Δ*frzE*, blue). Boxes extend from the 25th to 75th percentile, whiskers show 1.5× the interquartile range, and dots represent individual cells. Sample sizes across the six calcium conditions were: A⁺S⁺ (4 replicates): 36, 140, 249, 167, 271, and 164 cells; A⁺S⁻ (3 replicates): 19, 32, 31, 61, 45, and 34 cells; A⁻S⁺ (3 replicates): –, –, 29, 67, 147, and 199 cells.

Remarkably, A⁺S⁺ cells combined both behaviors: they glided with no detectable pili activity signature at 0 mM [CaCl₂], acquired leading-pole fluctuations from 0.4 mM onward, and at ≥0.9 mM the lagging pole progressively lost alignment with the cell center, indicating that the entire cell increasingly follows pilus-driven fluctuations initiated at the leading pole (Fig. 3c, Movie S11). Together, these observations suggest that wild-type cells gradually shift from adhesion-dominated to pili-dominated propulsion as calcium concentration increases, rather than relying on one system exclusively.

To better capture how motility patterns change with calcium, we summarized the 48 motility features into a two-dimensional “behavioral space” using UMAP (Fig. 3d; Fig. S3a-h). In this space, proximity reflects similarity in movement patterns. A⁺S⁻ and A⁻S⁺ mutants each progressed smoothly through distinct regions as [CaCl₂] increased, consistent with gradual modulation of a single motility system. By contrast, A⁺S⁺ cells occupied three separate regimes. At 0 mM [CaCl₂] they overlapped with the A⁺S⁻ cluster, consistent with pure gliding (Fig. 3d, left). At 4 mM [CaCl₂] they not only entered but extended beyond the A⁻S⁺ region, indicating amplified twitching dynamics (Fig. 3d, right). At intermediate concentrations (0.4-2 mM), A⁺S⁺ cells traced a clear transition zone: around 0.4 mM their profiles closely matched A⁺S⁻ cells at 0.9 mM, while already showing partial overlap with A⁻S⁺ cells at the same concentration; as [CaCl₂] rose toward 2 mM, they progressively entered the A⁻S⁺ region defined at higher calcium. This trajectory demonstrates that wild-type cells do not simply blend gliding and twitching behaviors but can also shift their dominant motility mode as calcium increases.

To assess the functional impact of these behavioral transitions, we next measured cell speeds across calcium concentrations. A⁺S⁻ cells maintained a rather constant speed profile peaking around 2mM [CaCl₂] (Fig. S3i), whereas A⁻S⁺ cells were immotile below ∼0.7 mM and then showed a sharp increase in velocity with rising calcium. These patterns confirm that calcium differentially modulates both motility systems: it subtly tunes adhesion-based gliding while strongly activating and amplifying pili-driven twitching. A⁺S⁺ cells combined features of both mutants: matching pure gliding speeds at zero calcium and ultimately surpassing each single-motor strain across the entire CaCl₂ range (Fig. 3e; Fig. S3i-k). Strikingly, at 4 mM [CaCl₂] the speed of A⁺S⁺ cells even outpaced the summed maximal speeds of the mutants, consistent with a synergistic interaction between A-and S-motility.

Together, these data indicate that calcium modulates each motility system individually but, more importantly, continuously shifts their relative contributions within the same cell, generating a continuum of propulsion regimes between gliding-and twitching-dominated movement. Notably, at high calcium the speed of wild-type cells exceeds the summed maxima of the single-motor strains, which is not expected if the two systems contributed independently.

### A-and S-motility systems couple non-additively

To test different hypotheses about how each motility system contributes to movement, we developed a biophysical simulation framework that models bacterial motility based on the underlying mechanisms. For this, we implemented an agent-based model in which the bacterium is represented as a deformable body, a linear chain of six disks connected by Hookean springs, with bending penalties to enforce shape persistence and velocity-proportional damping to capture surface friction and viscous drag (Methods) ^33–35^. This flexible architecture preserves overall cell integrity while allowing local deformations. Disk positions are updated using Verlet integration to ensure numerical stability. Importantly, all core mechanical parameters and nearly all motility-specific parameters (described hereafter) were drawn from the literature or directly constrained by our experimental data (Supplementary Table 3).

A-motility is implemented via focal adhesion complexes modeled as static substrate anchors that are deposited whenever the leading disk advances beyond a threshold distance from the previous anchor. Each focal adhesion complex generates a force aligned with the local axis of the cell and acts as a spring-like tether on any disk that overlaps with its position. This dual role captures both the directional propulsion and substrate anchoring characteristic of focal adhesion-based motility (Fig. 4a). To reflect intrinsic and cell-to-cell variability we varied key focal adhesion parameters (force magnitude, spring stiffness, deposition distance threshold) with noise within and between simulation runs. Because CaCl₂ affected A⁺S⁻ speed modestly and non-monotonically, we kept A-motility parameters CaCl₂-independent in the model rather than impose an unconstrained calcium dependence (Fig. 3e). This simulation produced straight, tightly aligned trajectories for the leading pole, center, and lagging pole (Fig. 4b, Movie S13), and instantaneous-speed distributions that resemble the experimental data across CaCl₂ levels (Fig. 4c).

**Figure 4.**
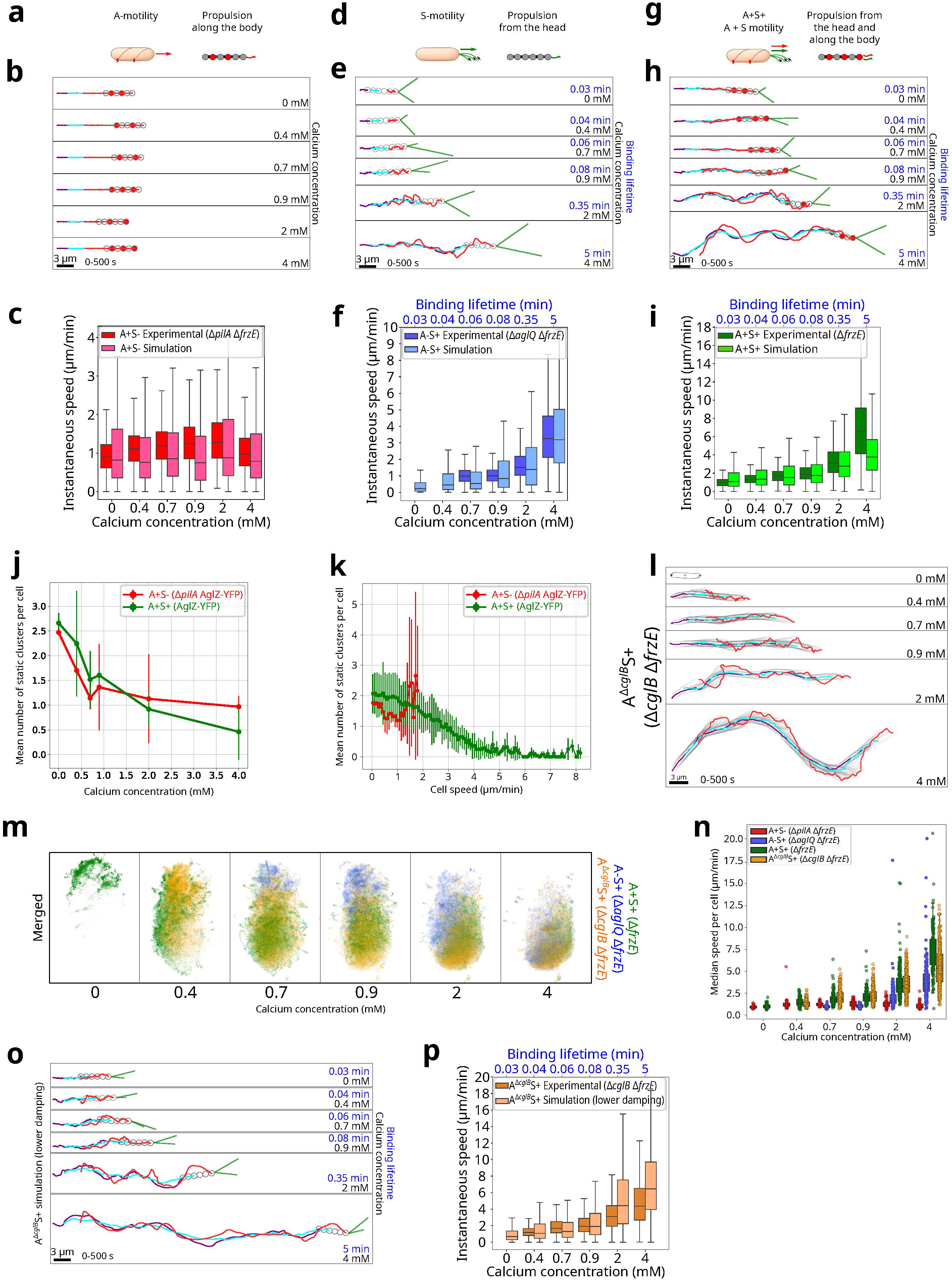
: Model Reproduces A⁺S⁻ and A⁻S⁺ Behaviors and Supports a Non-Additive Role of A-Motility in A⁺S⁺. **(a-c)** Modeling gliding (A⁺S⁻) mutants. (a) Schematic of our A⁺S⁻ (Δ*pilA* Δ*frzE*) model: the cell is represented as six disks (grey) linked by springs, with a constant propulsive force along its length (red). (b) Examples of simulated trajectories of leading pole (red), lagging pole (purple), and cell center (cyan) used to compare with experimental data under increasing CaCl₂ in the following panels. Final disks configuration is outlined in gray. (c) Box plots of instantaneous speeds for experimental (red) versus simulated (pink) A⁺S⁻ cells across CaCl₂ (n = 100 simulated trajectories per condition). Boxes: 25th-75th percentiles; whiskers: 1.5× interquartile range. **(d-f)** Modeling twitching (A⁻S⁺) mutants. (d) Schematic of our A⁻S⁺ (Δ*aglQ* Δ*frzE*) model: six disks with propulsion on the leading disk only, driven by pili whose binding lifetime varies with CaCl₂. (e) Examples of simulated trajectories for A⁻S⁺ driven by pili (green lines) under the same CaCl₂ series, where each concentration corresponds to a different binding lifetime. (f) Instantaneous speed box plots for experimental (blue) and simulated (light blue) A⁻S⁺ cells across CaCl₂/binding lifetimes (n = 100 simulated trajectories per condition). **(g-i)** Modeling A⁺S⁺. (g) Schematic of the combined A⁺S⁺ (Δ*frzE*) additive model, merging A⁺S⁻ and A⁻S⁺ modules with the same parameter sets. (h) Simulated A⁺S⁺ trajectories across CaCl₂/binding lifetimes. (i) Instantaneous speed box plots for experimental (green) and simulated (light green) A⁺S⁺ across CaCl₂/binding lifetimes (n = 100 simulated trajectories per condition). **(j)** Mean number of static AglZ-YFP clusters per cell across calcium concentration for experimental A⁺S⁻ (Δ*pilA*, red) and A⁺S⁺ (green) strains. Error bars = ±SD across replicates. Sample sizes across the six calcium conditions were: A⁺S⁻ (2 replicates): 13, 23, 24, 28, 16, and 17 cells; A⁺S⁺ (2 replicates): 7, 108, 127, 123, 91, and 53 cells. **(k)** Mean static AglZ-YFP clusters count versus instantaneous cell speed for the same strains (A⁺S⁻ n = 122; A⁺S⁺ n = 509, 2 replicates each). Error bars = ±SD across replicates. **(l-n)** Experimental results in a Δ*cglB* background. (l) Trajectories of the leading pole (red), lagging pole (purple), and center (cyan) for representative A^Δ*cglB*^S⁺ (Δ*cglB* Δ*frzE*) cells across calcium concentrations. Cell contours are shown at each timepoint (light gray). (m) UMAP density maps for experimental A⁻S⁺ (Δ*aglQ* Δ*frzE*, blue), A⁺S⁺ (Δ*frzE,* green), and A^Δ*cglB*^S⁺ (Δ*cglB* Δ*frzE*, orange), each pooled over CaCl₂ (0-4 mM). (n) Median instantaneous speeds per cell versus CaCl₂ for A⁺S⁺ (Δ*frzE,* green), A⁺S⁻ (Δ*frzE,* red), A⁻S⁺ (Δ*aglQ* Δ*frzE*, blue), and A^Δ*cglB*^S⁺ (Δ*cglB* Δ*frzE,* orange). Sample sizes for A⁺S⁺, and A⁻S⁺ are the same as in Figure 3e. Sample size for A^Δ*cglB*^S⁺ across the six calcium conditions were (N=4 replicates) –, 155, 253, 245, 342, and 390 cells. **(o-p)** Reduced-damping simulations. (o) Representative simulated A^Δ*cglB*^S⁺ trajectories with reduced damping across CaCl₂/binding lifetimes. (p) Instantaneous speed box plots for experimental A^Δ*cglB*^S⁺ (orange) and A^Δ*cglB*^S⁺ low-damping simulations (peach) across CaCl₂/binding lifetimes, n = 100 simulated trajectories per condition.

We next modeled A⁻S⁺ cells by modeling type IV pili activity at the leading pole (Fig. 4d). In this model, pili stochastically bind the substrate within a defined angular range and retract at 0.5 µm/s velocity ^19^. In a similar way to A⁺S⁻ modeling, we varied key pili parameters (binding rate, retraction rate, maximum pilus count, length, stall force, angular binding range) with noise within and between simulation runs to reflect intrinsic and cell-to-cell variability. In these simulations, we did not consider variations in pilus length or number with [CaCl_2_] as fluorescent labeling of pili revealed no consistent change in pilus length or number across calcium concentrations (Fig. S4a). Instead, we varied pilus binding lifetime as a proxy for calcium-mediated stabilization. Simulated trajectories closely resembled experimental patterns (Fig. 3b), with characteristic center-lagging pole alignment and enhanced leading pole fluctuations (Fig. 4e, Movie S14). Moreover, instantaneous speed distributions closely matched the experimental data (Fig. 4f). The only exception was below 0.4 mM CaCl₂, where experimental A⁻S⁺ cells were largely immobile while simulated cells remained motile, suggesting a potential biological activation threshold or regulatory mechanism that limits S-motility under low-calcium conditions in vivo, possibly linked to calcium-binding affinity.

We then asked whether simulations combining A⁺S⁻ and A⁻S⁺ models could recapitulate experimental A⁺S⁺ motility patterns (Fig. 4g). To address this question, we first constructed an “additive” A⁺S⁺ model without synergy (where both systems acted independently from each other). The simulated trajectories resembled those of experimental A⁺S⁺ cells at 2-4 mM CaCl₂ (Fig. 4g), with similar leading pole fluctuations and center-lagging pole coordination (Fig. 4h vs. 3c, Movie S15). However, below 0.9 mM the simulated cells showed reduced leading-pole fluctuations compared to the experiments, indicating they lack the dynamic leading pole deflections of true A⁺S⁺ cells. Next, we compared instantaneous speeds. Across most CaCl₂ levels (0-2 mM), the additive model tracked the experimental speed curves faithfully (Fig. 4i). Yet at 4 mM CaCl₂ the simulated cells showed lower speeds than experiments. Together, these results show that, within the confines of our simplified model, simply summing A-and S-motility without synergy fails to account for both leading pole fluctuation dynamics and velocities across the whole CaCl₂ range. Together, these results indicate that an additive model is insufficient to explain the experimental observations, supporting a non-additive interaction between the two motility systems.

### A-Motility Enhances S-Motility Independently of Focal Adhesions

To explore whether the two motility systems influence one another beyond simple co-deployment, we quantified focal adhesion complexes (FACS) across CaCl₂ levels. In A⁺S⁻ cells, the mean number of FACS decreased with rising [CaCl₂] and stabilized at roughly one adhesion per cell at high calcium (Fig. 4j; Fig. S4b,d, Movie S16), consistent with A-motility being the sole propulsion mechanism in this background and indicating that CaCl₂ also impacts the assembly of focal adhesions. In A⁺S⁺ cells, the mean number of FACS complexes per cell displayed a similar decrease with [CaCl_2_] below 0.9 mM. In contrast, the mean number of FACS was lower than one per cell above 2 mM (Fig. 4j; Fig. S4c,e, Movie S17). This is a surprising observation given that A⁺S⁺ speeds continued to increase with [CaCl_2_] above 2mM. These observations can be explained by a considerable proportion of A⁺S⁺ cells moving without detectable adhesions at high CaCl₂ concentrations. To test this hypothesis, we analyzed how the mean number of FACS varies with instantaneous cell speed. In A⁺S⁻ cells, the mean number of FACS remains above one up to the maximum speed of the cells (∼ 2 µm/min) (Fig. 4k). In contrast, in A⁺S⁺ cells the mean number of FACS per cell decreased steadily with speed, and on average, vanished above ∼6 µm/min (Fig. 4k, Fig. S4f-g). These observations demonstrate that, at high CaCl₂ levels, wild-type *M. xanthus* cells can achieve rapid, focal adhesion-independent movement. Yet A⁺S⁺ cells still outpace A⁻S⁺ mutants under the same conditions, suggesting that the A-system enhances S-motility through a mechanism other than direct traction by focal adhesions.

To test this directly, we examined an A-motility mutant lacking the surface adhesin but retaining a complete A-motor complex (Δ*cglB* Δ*frzE,* hereafter A^Δ*cglB*^S^+^). This strain thus retains pili-based propulsion and a non-anchored A-system with a functional A-motor activity. Kymographs of AglZ-YFP in A^Δ*cglB*^S^+^ cells confirmed the presence of dynamic A-motility complexes without any static adhesions (Fig. S4h, Movie S19). In absence of CaCl₂, A^Δ*cglB*^S^+^ cells were immotile, consistent with the absence of functional A-and S-motilities in this condition (Fig. 4l, Movie S18). In contrast, above 0.4 mM [CaCl₂], A^Δ*cglB*^S^+^ cells displayed increasing leading-pole oscillations and lagging-pole decoupling, closely mirroring the CaCl₂ response of A⁺S⁺ cells. Strikingly, at 0.4 mM [CaCl₂] A^Δ*cglB*^S^+^ cells no longer occupy the gliding-associated region seen in A⁺S⁺ cells but instead map onto the twitching-like sector characteristic of A⁻S⁺ behavior, indicating early pili engagement before the A⁻S⁺ mutant even moves, suggesting that the A-system facilitates or accelerates the onset of pili-based motility. At calcium levels above 0.7 mM, A^Δ*cglB*^S^+^ trajectories overlapped extensively with A⁺S⁺ cells, yet remained distinct from A⁻S⁺ cells (Fig. 4m; Fig. S4i-l). Speed measurements echoed this pattern: A^Δ*cglB*^S^+^ cells matched the velocities of A⁺S⁺ cells up to 2 mM and reached similar speeds at 4 mM (Fig. 4n; Fig. S4m). Notably, these findings show that the A-system can enhance S-motility even when CglB-mediated surface attachment is abolished, indicating that functional integration of the two systems does not require focal-adhesion-mediated traction.

Finally, we sought to rationalize how the A-system could facilitate S-motility without engaging substrate adhesion using simulations. First, we hypothesized that A^Δ*cglB*^S^+^ cells may have larger pili conferring them the ability to move faster than A⁻S⁺ cells. However, increasing the length of pili from 5 µm to 10 µm failed to reproduce the experimental speeds of A^Δ*cglB*^S^+^ cells, particularly underestimating instantaneous velocities at low CaCl₂ levels and remaining below experimental values even at high calcium concentrations (Fig. S4n). Second, we hypothesized that A^Δ*cglB*^S^+^ cells may have a higher number of pili. Doubling pilus number brought simulated velocities much closer to the data across the CaCl₂ concentration range, though it still underestimated speeds at intermediate CaCl₂ levels (Fig. S4o). Consistent with these in silico predictions, we observed no consistent differences in pilus length or number when we observed fluorescently labeled pili in A⁺S⁺ across a range of [CaCl_2_], from 0.7 to 4 mM (Fig. S4p). Third, we tested if variations in pilus retraction rate, maximal stall force, and mechanical damping with [CaCl_2_] can explain the behavior of A^Δ*cglB*^S^+^ cells. Simulations where we increased the retraction rate from 30 µm/min to 60 µm/min failed to improve the speed profiles, with simulated velocities consistently underestimated across all calcium concentrations (Fig. S4q). In contrast, either raising the stall-force limit (from 70 to 200 pN) or reducing the damping coefficient (from 8 to 2.5 pN·min/µm) yielded simulated trajectories that closely match the experimental patterns (Fig. 4o, Movie S20) and produced speed distributions that nearly superimpose on the experimental curves of A^Δ*cglB*^S^+^ cells across calcium concentrations (Fig. 4p; Fig. S4r). These results suggest that the A-motility system, even when unanchored, may facilitate S-motility by increasing pilus strength or reducing resistance to cellular motion.

## DISCUSSION

Many motile cells, from bacteria to eukaryotes, harbor more than one motility system, yet how mechanistically distinct systems are integrated within individual cells remains largely unknown. Here, we show that the evolutionarily distinct A-and S-motility systems of *M. xanthus* can be simultaneously active within single cells and jointly contribute to propulsion. These findings demonstrate that multiple motility systems need not function as alternative modes of locomotion, but can instead be integrated within the same cell to generate a unified and tunable mode of movement.

The observation that active A-motility operates in both isolated cells and large groups, together with the simultaneous use of A-and S-motility within individual cells, challenges the view that A-motility is dedicated to solitary movement whereas S-motility drives collective motility. In addition, the ability of single cells to use both motors expands the range of available cell behaviors: A-motility produces persistent, directional gliding suited for long-distance exploration, whereas S-motility promotes pole fluctuations, local steering, and cell-cell interactions ^36^. Engaging both allows single cells to move faster and explore space more effectively than cells relying on either system alone. This increased speed is observed on both chitosan-coated surfaces and agar, suggesting that this benefit is not specific to a single substrate. Consistently, wild-type swarms move faster than single-motility mutants, display higher directional persistence, and prey more efficiently ^26^, indicating that the advantage of dual-system use extends from single-cell exploration to collective predation. These findings raise a fundamental question: how are two evolutionarily and mechanistically distinct motility systems coordinated to produce coherent movement?

A shared polarity module provides a natural mechanism to coordinate the two motility systems. MglA and Frz control the polar assembly and reversals of both A-and S-motility systems ^22,23^, yet why these evolutionarily distinct systems are regulated by the same factors, and whether this shared regulation serves to coordinate their behavior has remained unclear. Our findings suggest that this shared regulation enables coordinated deployment of both motors within the same cell. In its GTP-bound form, MglA recruits the A-and S-machineries to the same pole through GltJ and SgmX, respectively ^20,21^, while Frz periodically switches this shared polarity axis during reversals. By polarizing both systems simultaneously, this common regulatory module can coordinate their activities while preventing a tug-of-war between opposite cell poles. Together, these observations suggest that the shared MglA/Frz polarity module determines where the two motors act, but not their relative contribution to propulsion. This raises the question of how the balance between A-and S-motility is regulated.

Our results identify extracellular calcium as an environmental signal that regulates the balance between the two motility systems. Across the environmentally relevant calcium range tested (0–4 mM CaCl₂, within the broader soil range of ∼0.01–14 mM ^37^), wild-type behavioral fingerprints shift progressively from A-motility-dominated to pilus-dominated propulsion, passing through co-propulsion at intermediate concentrations. Calcium concentration affects single-motility mutants differently, suggesting that the two systems can respond differently to the same environmental cue. The S-system carries a known calcium-sensing component, PilY1.1/minor-pilin complexes ^31,38^; the A-system carries a candidate calcium-sensing factor, the predicted MIDAS motif of CglB ^39^. Our simulations show that varying pilus binding lifetime reproduces the observed shift in propulsion, consistent with calcium modulating pilus-substrate binding as might occur through PilY1.1/minor-pilin complexes ^31^. Together, these findings suggest that a single environmental cue can continuously tune the relative output of two evolutionarily distinct motility systems through distinct molecular mechanisms in each motor, thereby shifting propulsion from gliding-to twitching-dominated regimes in response to changing external conditions.

Surprisingly, however, calcium-mediated tuning alone cannot explain the enhanced performance of wild-type cells, indicating that A-and S-motility interact non-additively rather than acting as fully independent motors. This conclusion is supported by several independent observations. First, at high calcium concentrations, wild-type cells reach speeds exceeding the summed maximal speeds of the two single-system mutants, and this supra-additivity cannot be reproduced by simulations in which A-and S-motility act independently. Moreover, wild-type speed is still enhanced despite the absence of stable focal adhesions in this regime, suggesting that the A-system contributes indirectly rather than through classical adhesion-based traction. Second, the ΔcglB mutant, whose A-motor remains functional but lacks stable focal adhesions, twitches faster than the A-motor-deficient mutant, indicating that this facilitation requires a functional A-motor rather than CglB-mediated adhesions. This enhancement is unlikely to result simply from the loss of CglB-dependent adhesive drag, since wild-type cells, which retain CglB, also move faster than the A-motor-deficient mutant. Among the mechanisms tested in simulations, the behavior of ΔcglB mutants is best explained either by amplifying pilus-generated force or by reducing mechanical resistance to motion. One possible mechanism, consistent with helical-rotor and viscous-interaction models of A-motility ^40^, is that trafficking A-motility units along the cell body generate additional propulsive thrust or modify viscous coupling at the cell-surface interface. Determining the physical basis of this non-additive coupling, and whether it reflects viscous cell-surface interactions or another form of communication between the two motility systems, will be an important direction for future work.

Beyond *Myxococcus*, our findings suggest that the coordinated use of multiple motility systems may represent a general strategy for adaptive cell movement. Many organisms combine evolutionarily distinct locomotion modes, including flagella and pili in bacteria, crawling and swimming in protists, or lamellipodia and blebs in migrating animal cells ^4,41–44^. Whether these systems can be simultaneously engaged within the same cell, however, remains largely unknown. This question is particularly relevant because Agl-like gliding motors occur beyond myxobacteria ^45,46^, and type IV pilus systems, some with calcium-sensitive regulation, are found across diverse bacteria, with related pilus machinery also present in archaea ^8,27^. Together, these observations raise the possibility that the coordinated and environmentally tunable use of evolutionarily distinct motility systems is more widespread than currently appreciated. Although the molecular mechanisms will likely differ across organisms, our findings suggest that integrating distinct motility systems within a single cell may represent a general strategy for generating flexible and environmentally tunable propulsion.

## Data availability

Data generated in this study were deposited at the public repository 10.5281/zenodo.16265273.

## Code availability

Python code for the analysis of the data is available at the public repositories https://doi.org/10.5281/zenodo.16265273 and https://github.com/AntoineLeGall/2025_Mas_et_al.

## Acknowledgments

This project was funded by the French National Research Agency (grant ID ANR-23-CE13-0036) and the CBS2 doctoral school (A.M. scholarship). The CBS is a member of the France-BioImaging, a national infrastructure supported by the French National Research Agency (ANR-10-INBS-04-01). We further thank Emmanuel Margeat (CBS) for providing maleimide fluorophores for Type IV pili labeling, Laetitia My for her help in strain construction and Flora Honore (LCB) for sharing her advices for studying *M. xanthus* in Ibidi-chitosan microfluidics chambers. Claude (Anthropic) was used for assistance with language editing. ChatGPT (OpenAI) was used for assistance with language editing and code troubleshooting. All analyses, interpretations, and conclusions were performed and verified by the authors.

## Author Contributions

A.L.G., A.M., M.N., and T.M. conceived the study and the design. A.M. built new strains, implemented the experimental protocols, and acquired the data. A.L.G. and A.M. analyzed the data. A.L.G. built the microscope. A.L.G. and A.M. wrote the software. A.L.G. and A.M. performed simulations. A.L.G., A.M., and M.N. interpreted the data. A.L.G. and A.M. wrote the first manuscript draft. A.L.G., A.M., M.N., and T.M. contributed to manuscript editing. A.L.G., M.N., and T.M. supervised the study and acquired funding.

## METHODS

### Predation assay

*M. xanthus* strains were grown in CYE (10 mM MOPS pH 7.6; 10 g/L Bacto Casitone peptone; 5 g/L yeast extract; 1 g/L MgSO₄·7H₂O; q.s. to 1 L with Milli-Q water) medium at 32°C with shaking (0.42 x g) overnight in the dark. Cells were centrifuged at 2 500 × g for 5 min, the supernatant discarded, and cells washed twice in CF buffer (10 mM MOPS pH 7.6; 1 mM KH₂PO₄; 8 mM MgSO₄; 0.02 % (NH₄)₂SO₄; 0.2 % sodium citrate; 0.015 % casitone, q.s. to 1 L with Milli-Q water) before being resuspended in CF buffer to an OD₆₀₀ of 5. *E. coli* prey cells were grown overnight in LB at 37 °C with shaking (0.42 x g) in the dark. The culture was then diluted 1:10 into fresh LB, incubated for 2 h, and centrifuged for 5 min at 2500 x g. The supernatant was discarded, and cells were resuspended in CF buffer to an OD₆₀₀ of 0.005. CF agar pads (0.75 % or 1.5 %) were prepared on microscope slides. On each pad, 1 µL of *M. xanthus* suspension was spotted, and 1 µL of *E. coli* suspension was spotted ∼0.5 mm away. Pads were then placed onto the surface of CF agar (1.5 %) in standard Petri dishes to maintain humidity, and incubated for 48 h at 32 °C. For imaging, slides were carefully removed, a coverslip was applied, and predation was observed by microscopy.

### Microfluidic chitosan device

*M. xanthus* cells were grown overnight in CYE medium at 32 °C with shaking (0.42 × g) in the dark. Cultures were centrifuged for 5 min at 2 500 × g, the supernatant discarded, and cells resuspended in CF buffer to an OD₆₀₀ of 0.05. The microfluidic device comprised an Ibidi sticky-slide (Ibidi VI 0.4) assembled with a chitosan-coated coverslip (Idylle). Before use, the chitosan surface was rehydrated by incubating with Milli-Q water for 15 min. To prepare the channel, it was first flushed once with *M. xanthus* suspension to remove any residual fluid, and then a second flush (90 µL) was used to establish the final cell suspension for imaging. To promote firm cell adhesion to the chitosan surface, the assembled slide was centrifuged for 1 min at 50 × g. For calcium-gradient experiments, the 90 µL of culture used for the final suspension was first removed, leaving the adhered cells on the substrate. Then, CF buffer supplemented with CaCl₂ (90 µL) was introduced sequentially into the same channel at final concentrations of 0.4, 0.7, 0.9, 2.0, and 4.0 mM. Each CF CaCl₂ solution was incubated for 2 min before imaging; the solution was then removed and replaced with the next concentration.

### Type-IV pili labelling and visualization

Pili were labeled using an adapted protocol based on ^21,47^. Strains of *M. xanthus* carrying pSWU19*-P_pilA_-pilA^D71C^* or pSWU30*-P_pilA_-pilA^D71C^*(see Supplementary Table 1) were grown overnight in CYE at 32 °C with shaking (0.42 × g) in the dark. Cultures were centrifuged (2 min at 4 930 × g), the supernatant discarded, and cells resuspended at an OD₆₀₀ of 0.3 in CF buffer containing CaCl₂ at the desired concentration and 25 µg.mL⁻¹ of Atto 565-maleimide (Atto-tec) or Alexa Fluor^TM^ 488 C_5_ maleimide (Thermo Fisher). Labeling proceeded in the dark for 1 h at room temperature. After incubation, cells were centrifuged (2 min at 4 830 × g) and washed seven times in CF CaCl₂ (same calcium concentration) to remove unbound fluorophores. Finally, cells were resuspended in CF CaCl₂ to an OD₆₀₀ of 0.1. For microfluidic experiments, chitosan-coated channels were rehydrated for 15 min with Milli-Q water and rinsed 2 times with CF CaCl₂. Labeled cells were introduced into the channels and centrifuged (1 min at 50 × g) before imaging. For imaging, time-lapse acquisitions were performed every 2 or 5 s with an exposure time of 10 ms, using either the 561 nm laser to image Atto 565 or the 488 nm laser for Alexa Fluor 488, in HILO illumination mode. Image processing was carried out using FIJI (ImageJ), including application of an unsharp mask to enhance fine structures and background subtraction to improve signal-to-noise ratio.

### Strains construction

For construction of AM1 (DZ2 att_Mx8_::pSWU30*-P_pilA_-pilA^D71C^* AglZ-YFP) and AM8 (DZ2 Δ*aglQ* att_Mx8_::pSWU30*-P_pilA_-pilA^D71C^*), a pSWU30-based plasmid encoding the *P_pilA_-pilA^D71C^* variant was integrated at the Mx8 phage attB site in TM9 (DZ2 *aglZ-YFP*) and TM146 (DZ2 Δ*aglQ*), respectively, via site-specific recombination. Plasmids were introduced by electroporation, and transformants were selected on CYE medium containing 5 µg/mL tetracycline. Correct insertion was confirmed by whole-genome sequencing (Plasmidsaurus) of DNA prepared with the Quick-DNA Miniprep Plus Kit (Zymo Research). Phenotypic motility assays were then performed on CYE agar (0.5% and 1.5%) by spotting 10 µL of OD₆₀₀ = 5 cultures and incubating for 48 h, using wild-type strain and known motility mutants as controls. Finally, the presence of pili was verified by microscopy.

### High-speed acquisition

Imaging was performed on a custom-built microscope based on a Nikon Eclipse Ti inverted stand, using a Nikon CFI Plan Apochromat Lambda 100×/1.45 NA oil-immersion objective (Nikon Instruments). For phase-contrast imaging, a ring-shaped LED illuminator (Amazon) was used in place of a traditional condenser to illuminate the sample from above. A PH3 phase ring, mounted in a plane conjugate to the objective’s back focal plane, was aligned to overlap with the LED ring image to generate phase contrast images on an Andor iXon Ultra 897 EMCCD (Andor Technology). For fluorescence excitation, two laser lines, 488 nm (Coherent CUBE) and 561 nm (Coherent Sapphire), were combined via dichroic mirrors and their intensities computer-controlled by an acousto-optic tunable filter (AOTF). The selected beam was then directed through a dichroic set FF496-SDi01 (Semrock) to switch between total internal reflection fluorescence (TIRF) and highly inclined laminated optical sheet (HILO) excitation paths. Fluorescence signal was then separated from excitation light using a dichroic ZT 405/488/561/633rpc (Chroma Technology) and separated from LED illumination channel using a second dichroic FF484-FDi01 (Semrock) transmitting fluorescence signal onto a second Andor iXon Ultra 897 EMCCD camera. Emitted light was filtered through a 525/50 nm band-pass for YFP/Alexa 488 or a 605/70 nm band-pass for mCherry/Atto 565 (Chroma Technology). The effective pixel size of both phase contrast and fluorescence cameras at the sample plane was 0.120 µm. All illumination pulses were controlled by a DAQ card and synchronized to the cameras integration windows: LED exposures were 400 ms, 488 nm laser pulses 10 or 30 ms, and 561 nm pulses 10 ms. Pulses were time-shifted to avoid overlap, and the three exposures cycled once per second to generate a complete illumination sequence of typically 16 minutes. An OBIS 785 nm infrared laser autofocus module (Coherent) maintained focus by monitoring back-reflections during acquisitions. All acquisitions were performed at room temperature (22-25 °C).

### Data preprocessing : Acquisition output, alignment, drift correction, and preprocessing

After microscope acquisition, a phase-contrast TIFF stack was generated, and zero, one, or two fluorescence stacks were produced depending on the experiment. When both 488 nm and 561 nm channels were acquired, they were stored in a single interleaved TIFF and then split into two separate stacks. When AglZ-YFP was imaged (488 nm channel), that fluorescence stack was denoised with the Aydin software ^48^ with the “Noise2SelfFGR-cb*”* model. Then each fluorescence channel was spatially aligned to the phase-contrast reference by estimating a pure translation via frequency-domain cross-correlation and applying this subpixel shift to the image stack, ensuring precise overlap between channels. To correct for any sample drift, the phase-contrast reference stack was temporally smoothed with a moving-average filter and intensity-clipped; OpenCV’s phase-correlation then measures frame-to-frame drift within a sliding window, and the inverse shifts were applied to each stack. Finally, the drift-corrected phase-contrast stack was processed frame by frame with an unsharp-mask filter to enhance edges.

### Segmentation

For each frame of the preprocessed phase-contrast stack, we ran the “bact_phase_omni” Cellpose model to generate a labeled instance mask that assigns every cell an integer label ^49^. The resulting per-frame masks were concatenated into a single TIFF stack, with each pixel’s value indicating the mask ID for that cell in that frame.

### Single-cell tracking

Single-cell tracks were reconstructed from the labeled phase-contrast TIFF stack using our custom Python pipeline (see Supplementary Data). Briefly, we tracked each cell’s centroid across frames using LapTrack’s cost-based mask matching ^50^, reconnected any fragmented trajectories to fill gaps, and discarded tracks that were too short or touched image borders. After finalizing track continuity, we relabelled each track with consecutive IDs and updated the labeled TIFF stack so that pixel values reflected these corrected labels. We then extracted per-cell morphological and kinetic features from each track’s masks, identified cell poles via skeleton-and-curvature analysis, and appended pole coordinates to the existing centroid tracks to obtain pole trajectories. Next, we smoothed all trajectories using a Savitzky-Golay filter to reduce high-frequency noise and computed signed displacements to distinguish leading versus lagging poles. Finally, we performed manual inspection in Napari ^51^ and applied morphological and kinetic filters to retain only high-quality single-cell trajectories for downstream analysis.

### Instantaneous speed calculation

Instantaneous speeds were computed directly on the non-rediscretized center trajectories by measuring the Euclidean distance between each pair of successive (x,y) positions and dividing by the acquisition interval (1 s), and converted to µm/min using the pixel size (0.120 µm pixel⁻¹).

### Trajectory discretization

To decouple spatial sampling from cell speed, we performed spatial trajectory rediscretization using Traja ^52^ to each center and pole track, placing new points at uniform 0.025-µm intervals.

### Cell features

For each spatially rediscretized cell trajectory, comprising its center, leading pole, and lagging pole, we computed a suite of 16 local morphodynamic features. Fifteen of these were derived via sliding fixed-size windows over the (x, y, time) data, and one (distances between coordinate pairs) was calculated instantaneously at each frame. Each feature quantified a specific aspect of cell shape or motion. Detailed implementation is provided in the Supplementary Data.

### UMAP embedding

UMAP embedding was computed on the preprocessed 48-dimensional feature matrix (see Supplementary Data for preprocessing procedure), comprising only our local morphodynamic descriptors (excluding dynamic features such as instantaneous speed), using the umap-learn package with the following settings: n_neighbors=50, min_dist=0.9, metric=“euclidean”, init=“spectral”, n_components=2. The resulting two-dimensional layout was rendered with Datashader, color-coding points by strain or calcium concentration, to reveal phenotype clusters across conditions. To compare global distributions, we overlaid iso-density contours computed via Gaussian kernel density estimation on a 2D grid, allowing visual comparison of population structure across conditions.

### Polar density plot

We visualized the relationship between angular deviation and their corresponding speeds by constructing a polar density plot. First, we selected the leading pole angular deviations (in degrees) and leading pole speeds (µm/min) for the strain and condition of interest, then determined a radial limit (rmax) as the 95th percentile of all speeds to avoid distortion by extreme outliers. We binned the data into a two-dimensional histogram over a regular grid of angles (−180° to +180°) and speeds (0 to rmax), applied a small Gaussian filter to smooth the resulting counts, and plotted the smoothed density in polar coordinates. In this view, each direction around the circle corresponds to a turning angle (0° at top, increasing clockwise to the right and counterclockwise to the left), and the distance from the center encodes speed. High-density regions thus reveal the predominant movement directions and speeds of the leading pole under the chosen experimental conditions.

### Automated kymographs generation

For experiments imaging AglZ-YFP, time-lapse sequences were acquired every 1-5 s with a 30 ms exposure. Kymographs were computed on the fluorescence TIFF stack that had been drift-corrected, denoised, and rigidly registered to the phase-contrast channel. Because cells often bend, we computed their medial axis for each one of them in each frame by skeletonizing their segmented masks and fitting that skeleton with a smooth spline to define the long axis. We then sampled fluorescence intensity along each cell length by, at each spline point, drawing a short line segment of fixed width perpendicular to the spline and averaging the pixel values along it, producing a raw kymograph (intensity vs. position) for each timepoint. Cell motion correction was then applied by projecting each frame-to-frame cell-centroid displacement onto the long-axis direction and shifting the corresponding kymograph profile by that amount, thereby fixing the kymograph in the substrate reference frame so that fluorescence features stationary on the surface remain static over time.

### Clusters of AglZ-YFP classification

AglZ clusters were first detected using the Spotiflow probabilistic spot detector and refined by 2D Gaussian fitting ^53^.These detections were then linked into tracks with TrackPy ^54^, gap-filled by linear interpolation, and tracks shorter than five frames were discarded. The remaining trajectories were smoothed with a Savitzky-Golay filter, spatially rediscretized, and a suite of morphodynamic features was computed. We then used a seven-dimensional Gaussian Mixture Model to classify every spot-timepoint as “static” or “mobile.” Finally, we assigned each cluster track to its nearest cell mask and retained only those clusters located more than 5 px away from the leading pole, thereby selecting for adhesion complexes that move along the cell’s basal surface within the substrate-proximal zone for downstream analysis.

### Simulations

To model bacterial motility, we implemented a two-dimensional agent-based simulation in which each cell was represented as a chain of six interconnected disks. Disks were linked by Hookean springs and subjected to bending and damping forces to capture the mechanical properties of the cell body. Bending penalties enforced shape persistence, while damping simulated resistance from surface friction and viscous drag. Disk positions were updated using Verlet integration to ensure numerical stability. Two propulsion systems were modeled: type IV pili and focal adhesion complexes (FACS). Pili extended from the leading disk and bound stochastically to the substrate within a defined angular range. Retraction generated pulling forces limited by a motor saturation threshold. Binding durations were randomly sampled around a user-defined lifetime, independent of force feedback. FACS were modeled as transient, static adhesive sites deposited when the leading disk advanced beyond a threshold distance from the previous anchor. Each FACS acted as a spring-like tether that exerted a restoring force on any overlapping disk and simultaneously applied a directional force aligned with the local cell axis, defined by the vector from disk *i-1* to *i+1*. To capture biological heterogeneity, core pili parameters (e.g., rest length, binding rate, retraction rate, number, motor force, angular range), and FACS parameters (e.g., force magnitude, spring stiffness, anchoring threshold) were independently perturbed using multiplicative log-normal noise (σ = 50%) in each simulation. Full simulation and processing details are provided in the Supplementary Methods.

