## Supplementary Information for "Integration of evolutionarily distinct motility systems enables tunable cell propulsion"

### Figure S1

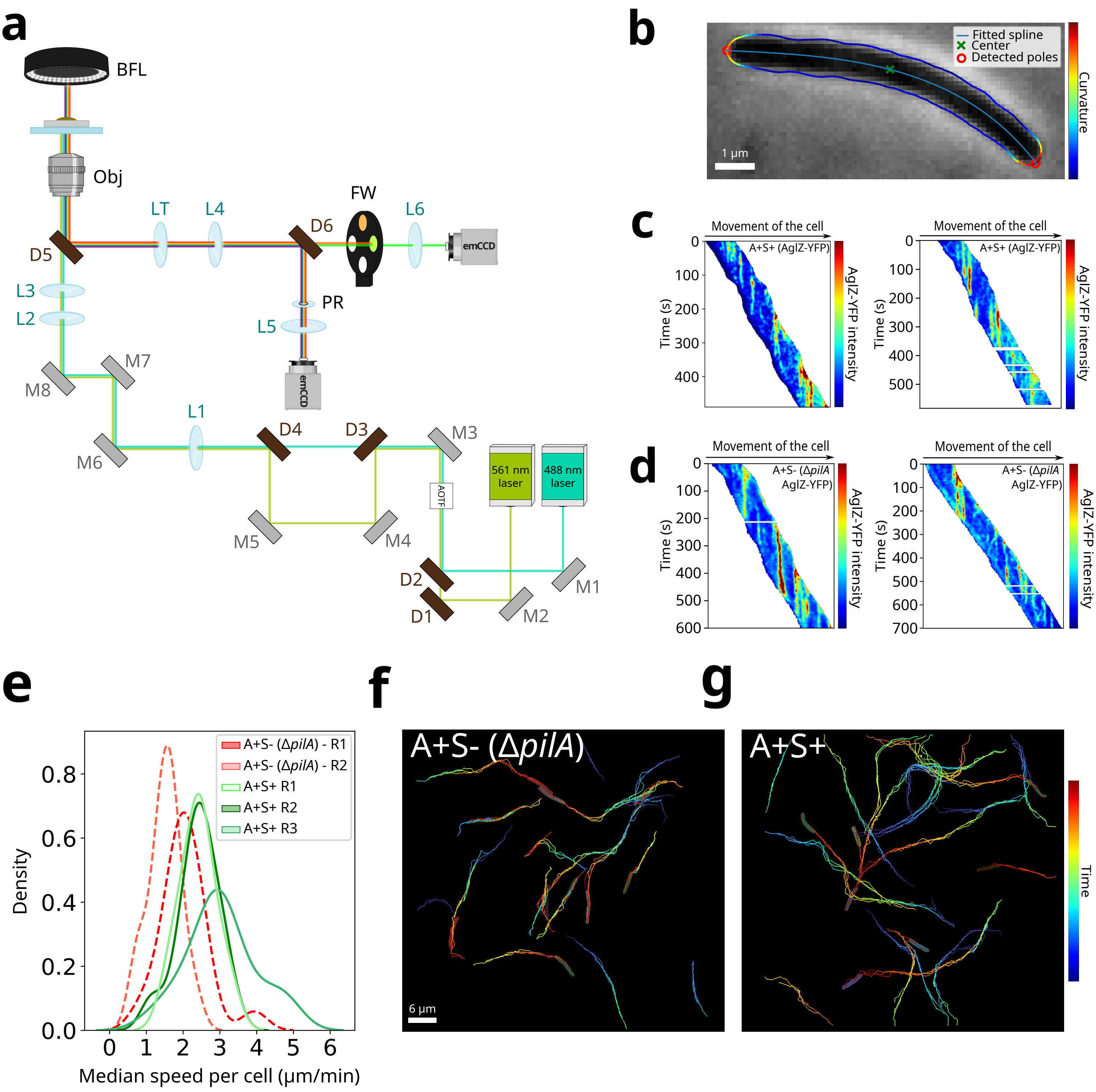

a
BFL
Obj
LT
L4
FW
L6
D6
D5
emCCD
L3
L2
PR
L5
M8
M7
L1
D4
D3
M3
M6
561 nm laser
488 nm laser
AOTF
M5
M4
D2
M1
D1
M2
b
Fitted spline
Center
Detected poles
Curvature
1 μm
c
Movement of the cell
A+S+ (AglZ-YFP)
Time (s)
AglZ-YFP intensity
d
A+S- (ΔpilA AglZ-YFP)
e
A+S- (ΔpilA) - R1
A+S- (ΔpilA) - R2
A+S+ R1
A+S+ R2
A+S+ R3
Density
Median speed per cell (μm/min)
f
A+S- (ΔpilA)
6 μm
g
A+S+
Time

### Supplementary Figure 1

(a) Schematic of the microscope setup used for high-speed sequential acquisition, enabling alternating TIRF (488 nm) and HILO (561 nm) fluorescence imaging, coupled with phase contrast acquisition. Excitation light from two lasers is directed to the sample through a 100x objective (Obj) via a sequence of mirrors (M1-M8), lenses (L1-L6), and dichroic mirrors (D1-D6), enabling independent tuning of TIRF and HILO modes. Emitted fluorescence is filtered through emission filters positioned within a filter wheel (FW) and detected on an EMCCD camera. Phase-contrast illumination was generated using a ring-shaped LED bright-field light (BFL) illuminating the sample from above. Transmitted light was collected by the objective, passed through a PH3 phase ring (PR), mounted in a plane conjugate to the objective's back focal plane, and imaged onto an EMCCD camera. The geometry and distances are illustrative and not drawn to scale.
(b) Pole and center detection pipeline. A spline is fitted to the cell's skeleton (light blue line), then extended to intersect the cell contour (blue outline). Local curvature along the contour is computed and color-coded from low (blue) to high (red). The cell poles are identified as points of maximum curvature and are shown as red circles. The geometric center of the cell is marked with a green cross, corresponding to the midpoint along the spline.
(c-d) Automated kymographs of AglZ-YFP signal in $A^+S^+$ (c) and $A^+S^-$ (Δ*pilA*) (d) cells.
(e) Median instantaneous speeds distribution per cell for $A^+S^-$ (Δ*pilA*) and $A^+S^+$ strains across multiple replicates. $A^+S^-$ - R1 : n = 19 cells, R2 : n = 23 cells. $A^+S^+$ - R1 : n = 6 cells, R2 : n = 26 cells, R3 : n = 22 cells.
(f-g) Trajectories of the leading pole, lagging pole, and center in $A^+S^-$ (Δ*pilA*) (f) and $A^+S^+$ (g) cells, color-coded by time over 1000 s.

### Figure S2

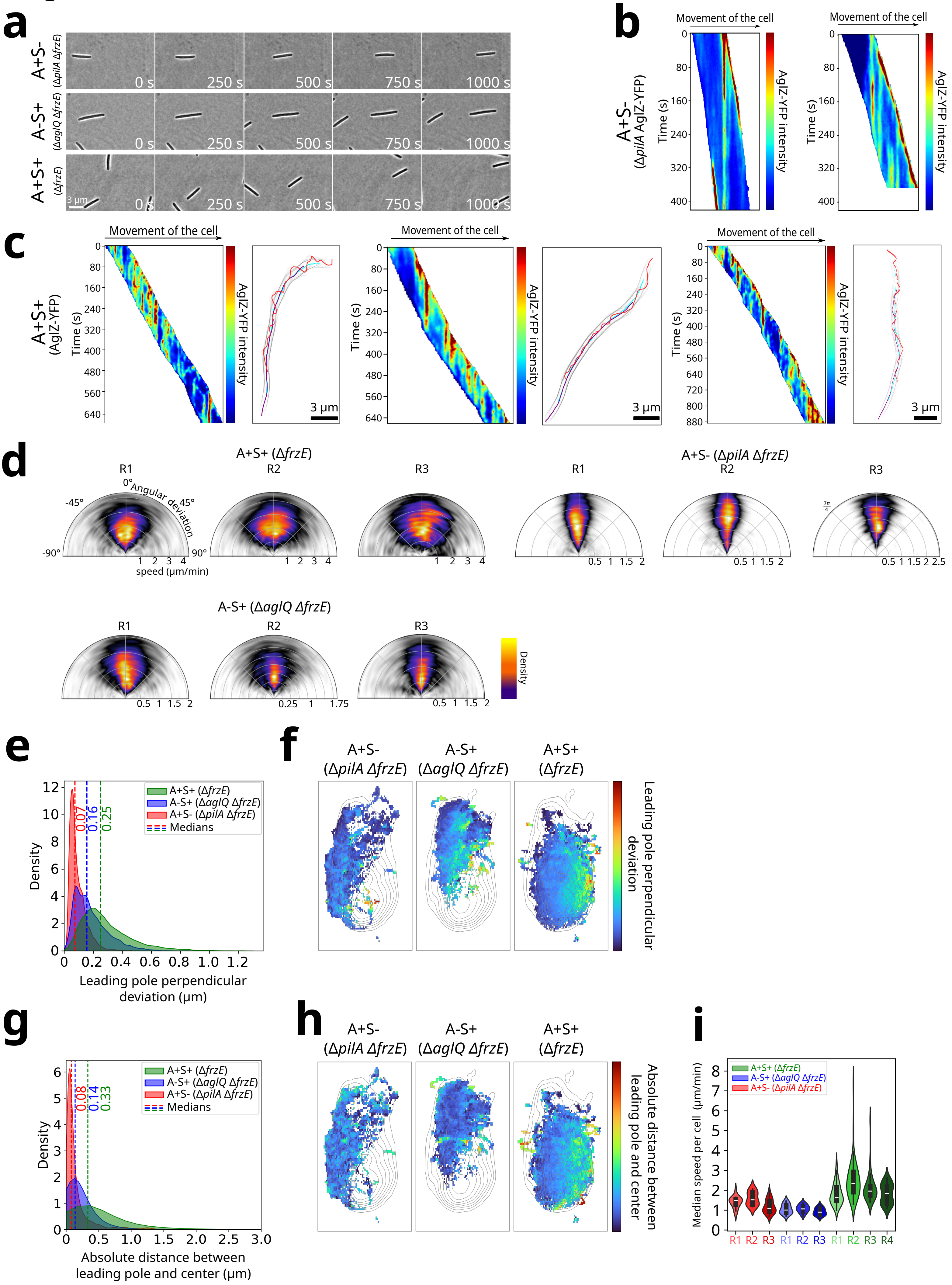
a
A+S- (ΔpilA ΔfrzE)
A-S+ (ΔaglQ ΔfrzE)
A+S+ (ΔfrzE)
0 s
250 s
500 s
750 s
1000 s
3 µm
b
A+S- (ΔpilA AglZ-YFP)
Movement of the cell
Time (s)
AglZ-YFP intensity
c
A+S+ (AglZ-YFP)
Movement of the cell
Time (s)
AglZ-YFP intensity
3 µm
d
A+S+ (ΔfrzE)
A+S- (ΔpilA ΔfrzE)
A-S+ (ΔaglQ ΔfrzE)
R1
R2
R3
Angular deviation
speed (µm/min)
Density
e
A+S+ (ΔfrzE)
A-S+ (ΔaglQ ΔfrzE)
A+S- (ΔpilA ΔfrzE)
Medians
0.07
0.16
0.25
Density
Leading pole perpendicular deviation (µm)
f
A+S- (ΔpilA ΔfrzE)
A-S+ (ΔaglQ ΔfrzE)
A+S+ (ΔfrzE)
Leading pole perpendicular deviation
g
A+S+ (ΔfrzE)
A-S+ (ΔaglQ ΔfrzE)
A+S- (ΔpilA ΔfrzE)
Medians
0.08
0.14
0.33
Density
Absolute distance between leading pole and center (µm)
h
A+S- (ΔpilA ΔfrzE)
A-S+ (ΔaglQ ΔfrzE)
A+S+ (ΔfrzE)
Absolute distance between leading pole and center
i
A+S+ (ΔfrzE)
A-S+ (ΔaglQ ΔfrzE)
A+S- (ΔpilA ΔfrzE)
Median speed per cell (µm/min)
R1
R2
R3
R4

### Supplementary Figure 2

**(a)** Phase contrast images of $A^+S^-$ (Δ*pilA* Δ*frzE*), A-S+ (Δ*aglQ* Δ*frzE)* and $A^+S^+$ (Δ*frzE*) on chitosan microfluidic device supplemented with 0.9 mM $CaCl_2$

**(b)** Automated kymographs of AglZ-YFP signal in $A^+S^-$ (Δ*pilA*) cells.

**(c)** Automated kymographs of AglZ-YFP signal in $A^+S^+$ cells, with corresponding tracking trajectories (leading pole in red, lagging pole in purple, center in cyan). Three examples are shown.

**(d)** Angular deviation (°) of the leading pole as a function of cell speed (µm/min) for three replicates (R1-3) of $A^+S^+$ (Δ*frzE*), $A^+S^-$ (Δ*pilA* Δ*frzE*), and $A^-S^+$ (Δ*aglQ* Δ*frzE*), color-coded by density. $A^+S^+$ - R1 : n = 38 cells, R2 : n = 44 cells, R3 : n = 26 cells. $A^+S^-$ - R1 : n = 33 cells, R2 : n = 18 cells, R3 : n = 10 cells. $A^-S^+$ - R1 : n = 27 cells, R2 : n = 20 cells, R3 : n = 20 cells.

**(e)** Distribution of the leading pole perpendicular deviation for $A^+S^+$ (Δ*frzE,* green, N = 4 replicates, n = 159 cells), $A^+S^-$ (Δ*pilA* Δ*frzE*, red, N = 3 replicates, n = 61 cells), and $A^-S^+$ (Δ*aglQ* Δ*frzE*, blue, N = 3 replicates, n = 67 cells) strains. Dashed line indicates the median value.

**(f)** UMAP projections of the full 48-dimensional behavior feature set for $A^+S^-$ (Δ*pilA* Δ*frzE*), $A^-S^+$ (Δ*aglQ* Δ*frzE*), and $A^+S^+$ (Δ*frzE*) cells. Rather than showing raw point density, each bin in the two-dimensional map is color-coded by the average normalized perpendicular deviation of the leading pole for all behavior segments falling into that bin. Iso-contours overlay the combined occupancy (density) across all three strains.

**(g)** Distribution of the absolute distance between leading pole and center for $A^+S^+$ (Δ*frzE,* green, N = 4 replicates, n = 159 cells), $A^+S^-$ (Δ*pilA* Δ*frzE*, red, N = 3 replicates, n = 61 cells), and $A^-S^+$ (Δ*aglQ* Δ*frzE,* blue, N = 3 replicates, n = 67 cells) strains. Dashed line indicates the median value.

**(h)** UMAP projections of the full 48-dimensional behavior feature set for $A^+S^-$ (Δ*pilA* Δ*frzE*), $A^-S^+$ (Δ*aglQ* Δ*frzE*), and $A^+S^+$ (Δ*frzE*) cells. Rather than showing raw point density, each bin in the two-dimensional map is color-coded by the absolute distance between leading pole and center for all behavior segments falling into that bin. Iso-contours overlay the combined occupancy (density) across all three strains.

**(i)** Median instantaneous speed per cell for $A^+S^+$ (Δ*frzE*), $A^+S^-$ (Δ*pilA* Δ*frzE*), and $A^-S^+$ (Δ*aglQ* Δ*frzE*) strains across all replicates. Dashed line indicates the median value. $A^+S^-$ - R1 : n= 18 cells, R2 : 10 cells, R3 : n = 33 cells. $A^-S^+$ - R1 : n= 20 cells, R2 : n= 27 cells, R3 : n= 20 cells. $A^+S^+$ - R1 : n= 40 cells, R2 : n= 54 cells, R3 : n= 46 cells, R4 : n= 27 cells.

### Figure S3

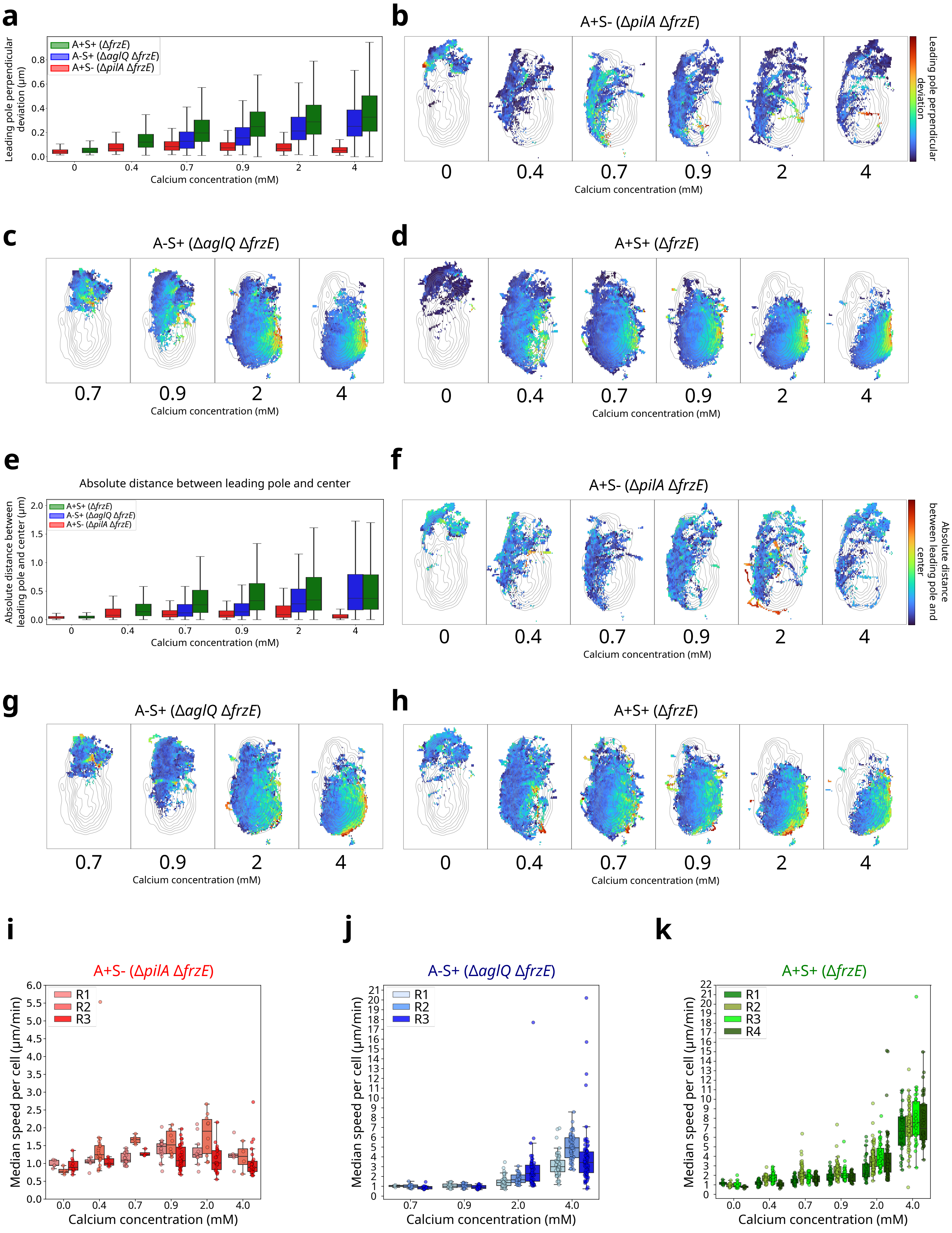

a
A+S+ (ΔfrzE)
A-S+ (ΔaglQ ΔfrzE)
A+S- (ΔpilA ΔfrzE)
Leading pole perpendicular deviation (µm)
Calcium concentration (mM)
b
A+S- (ΔpilA ΔfrzE)
Leading pole perpendicular deviation
0 0.4 0.7 0.9 2 4
Calcium concentration (mM)
c
A-S+ (ΔaglQ ΔfrzE)
0.7 0.9 2 4
Calcium concentration (mM)
d
A+S+ (ΔfrzE)
0 0.4 0.7 0.9 2 4
Calcium concentration (mM)
e
Absolute distance between leading pole and center
A+S+ (ΔfrzE)
A-S+ (ΔaglQ ΔfrzE)
A+S- (ΔpilA ΔfrzE)
Absolute distance between leading pole and center (µm)
Calcium concentration (mM)
f
A+S- (ΔpilA ΔfrzE)
Absolute distance between leading pole and center
0 0.4 0.7 0.9 2 4
Calcium concentration (mM)
g
A-S+ (ΔaglQ ΔfrzE)
0.7 0.9 2 4
Calcium concentration (mM)
h
A+S+ (ΔfrzE)
0 0.4 0.7 0.9 2 4
Calcium concentration (mM)
i
A+S- (ΔpilA ΔfrzE)
R1
R2
R3
Median speed per cell (µm/min)
0.0 0.4 0.7 0.9 2.0 4.0
Calcium concentration (mM)
j
A-S+ (ΔaglQ ΔfrzE)
R1
R2
R3
Median speed per cell (µm/min)
0.7 0.9 2.0 4.0
Calcium concentration (mM)
k
A+S+ (ΔfrzE)
R1
R2
R3
R4
Median speed per cell (µm/min)
0.0 0.4 0.7 0.9 2.0 4.0
Calcium concentration (mM)

### Supplementary Figure 3

**(a)** Distribution of the leading pole perpendicular deviation for $A^+S^+$ (Δ*frzE,* green), $A^+S^-$ (Δ*pilA* Δ*frzE*, red), and $A^-S^+$ (Δ*aglQ* Δ*frzE*, blue) strains across calcium concentration. $A^+S^+$ - N = 4 replicates, n = 36, 136, 244, 159, 263, 158 cells at 0, 0.4, 0.7, 0.9, 2, 4 mM. $A^+S^-$ - N = 3 replicates, n = 19, 32, 31, 61, 43, 31 cells at 0, 0.4, 0.7, 0.9, 2, 4 mM. $A^-S^+$ - N = 3 replicates, n = –, –, 29, 67, 145, 190 cells at 0, 0.4, 0.7, 0.9, 2, 4 mM.

**(b-d)** UMAP projections of the full 48-dimensional behavior feature set for $A^+S^-$ (Δ*pilA* Δ*frzE*) (b), $A^-S^+$ (Δ*aglQ* Δ*frzE*) (c), and $A^+S^+$ (Δ*frzE*) (d) cells across calcium concentration. Rather than showing raw point density, each bin in the two-dimensional map is color-coded by the average normalized perpendicular deviation of the leading pole for all behavior segments falling into that bin. Iso-contours overlay the combined occupancy (density) across all three strains and calcium concentrations.

**(e)** Distribution of the absolute distance between leading pole and center for $A^+S^+$ (Δ*frzE,* green), $A^+S^-$ (Δ*pilA* Δ*frzE*, red), and $A^-S^+$ (Δ*aglQ* Δ*frzE*, blue) strains. $A^+S^+$ - N = 4 replicates, n = 36, 136, 244, 159, 263, 158 cells at 0, 0.4, 0.7, 0.9, 2, 4 mM. $A^+S^-$ - N = 3 replicates, n = 19, 32, 31, 61, 43, 31 cells at 0, 0.4, 0.7, 0.9, 2, 4 mM. $A^-S^+$ - N = 3 replicates, n = –, –, 29, 67, 145, 190 cells at 0, 0.4, 0.7, 0.9, 2, 4 mM

**(f-h)** UMAP projections of the full 48-dimensional behavior feature set for $A^+S^-$ (Δ*pilA* Δ*frzE*) (f), $A^-S^+$ (Δ*aglQ* Δ*frzE*) (g), and $A^+S^+$ (Δ*frzE*) (h) cells across calcium concentration. Rather than showing raw point density, each bin in the two-dimensional map is color-coded by the absolute distance between leading pole and center for all behavior segments falling into that bin. Iso-contours represent the combined behavioral density of all three strains pooled at the calcium concentration of 0.9 mM.

**(i-k)** Median instantaneous speed per cell for strains across calcium concentrations and replicates. (i) $A^+S^-$ (Δ*pilA* Δ*frzE*). R1 : n = 4, 6, 22, 18, 11, 5 cells at 0, 0.4, 0.7, 0.9, 2, 4 mM. R2 : n = 5, 21, 5, 10, 10, 4 cells at 0, 0.4, 0.7, 0.9, 2, 4 mM. R3 : n = 10, 5, 4, 33, 24, 25 cells at 0, 0.4, 0.7, 0.9, 2, 4 mM. (j) $A^-S^+$ (Δ*aglQ* Δ*frzE*). R1 : n = –, –, 3, 20, 54, 67 cells at 0, 0.4, 0.7, 0.9, 2, 4 mM. R2 : n = –, –, 16, 27, 33, 54 cells at 0, 0.4, 0.7, 0.9, 2, 4 mM. R3 : n = –, –, 10, 20, 60, 78 cells at 0, 0.4, 0.7, 0.9, 2, 4 mM. (k) $A^+S^+$ strain (Δ*frzE*). R1 : n = 7, 51, 54, 40, 31, 30 cells at 0, 0.4, 0.7, 0.9, 2, 4 mM. R2 : n = 12, 32, 98, 54, 127, 62 cells at 0, 0.4, 0.7, 0.9, 2, 4 mM. R3 : n = 11, 42, 54, 46, 43, 34 cells at 0, 0.4, 0.7, 0.9, 2, 4 mM. R4 : n = 6, 15, 43, 27, 70, 38 cells at 0, 0.4, 0.7, 0.9, 2, 4 mM.

### Figure S4

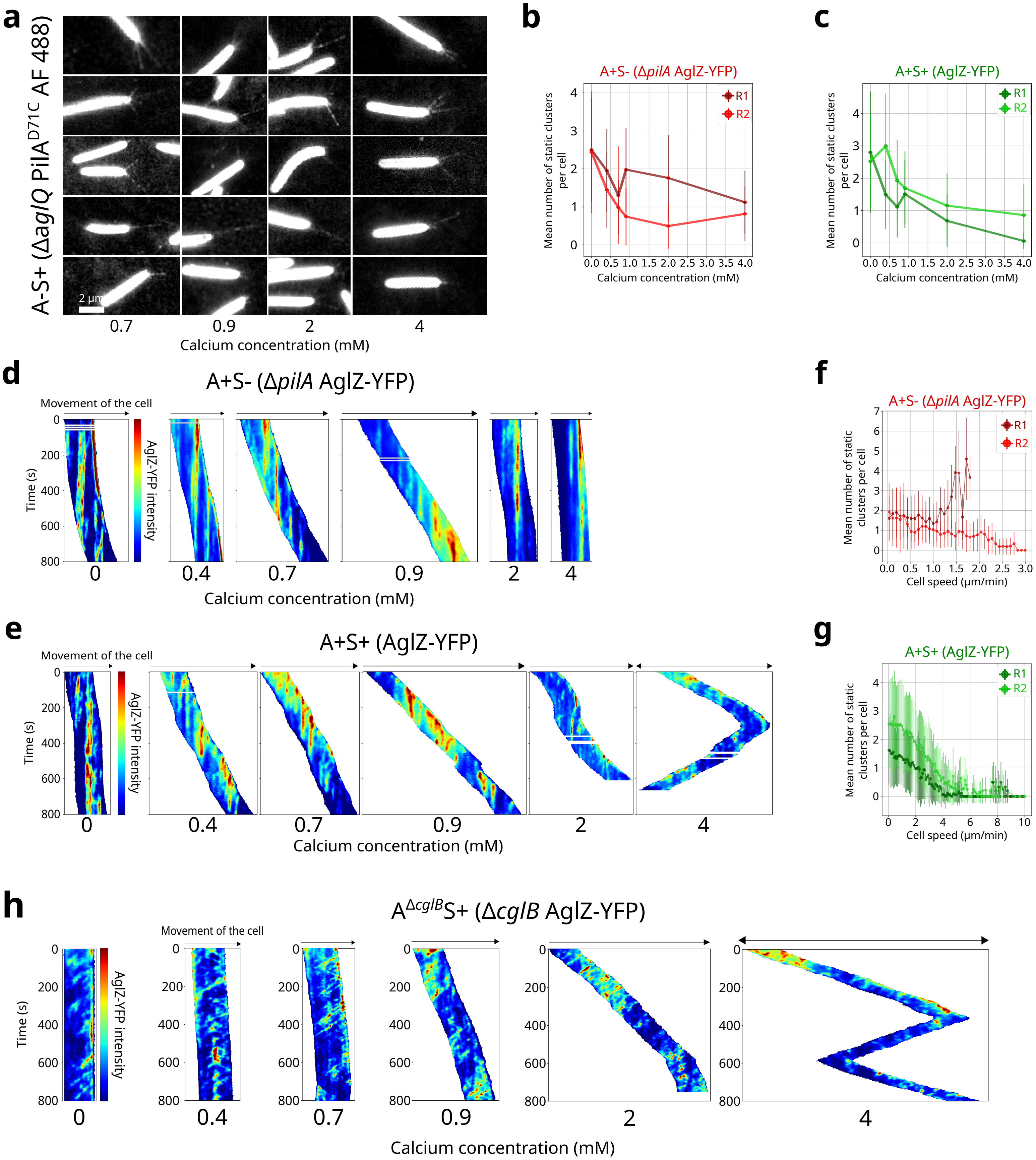
a
A-S+ (ΔaglQ PilA^D71C AF 488)
2 µm
0.7
0.9
2
4
Calcium concentration (mM)
b
A+S- (ΔpilA AglZ-YFP)
R1
R2
Mean number of static clusters per cell
Calcium concentration (mM)
c
A+S+ (AglZ-YFP)
R1
R2
Mean number of static clusters per cell
Calcium concentration (mM)
d
A+S- (ΔpilA AglZ-YFP)
Movement of the cell
Time (s)
AglZ-YFP intensity
0
0.4
0.7
0.9
2
4
Calcium concentration (mM)
f
A+S- (ΔpilA AglZ-YFP)
R1
R2
Mean number of static clusters per cell
Cell speed (µm/min)
e
A+S+ (AglZ-YFP)
Movement of the cell
Time (s)
AglZ-YFP intensity
0
0.4
0.7
0.9
2
4
Calcium concentration (mM)
g
A+S+ (AglZ-YFP)
R1
R2
Mean number of static clusters per cell
Cell speed (µm/min)
h
A^ΔcglB S+ (ΔcglB AglZ-YFP)
Movement of the cell
Time (s)
AglZ-YFP intensity
0
0.4
0.7
0.9
2
4
Calcium concentration (mM)

### Figure S4

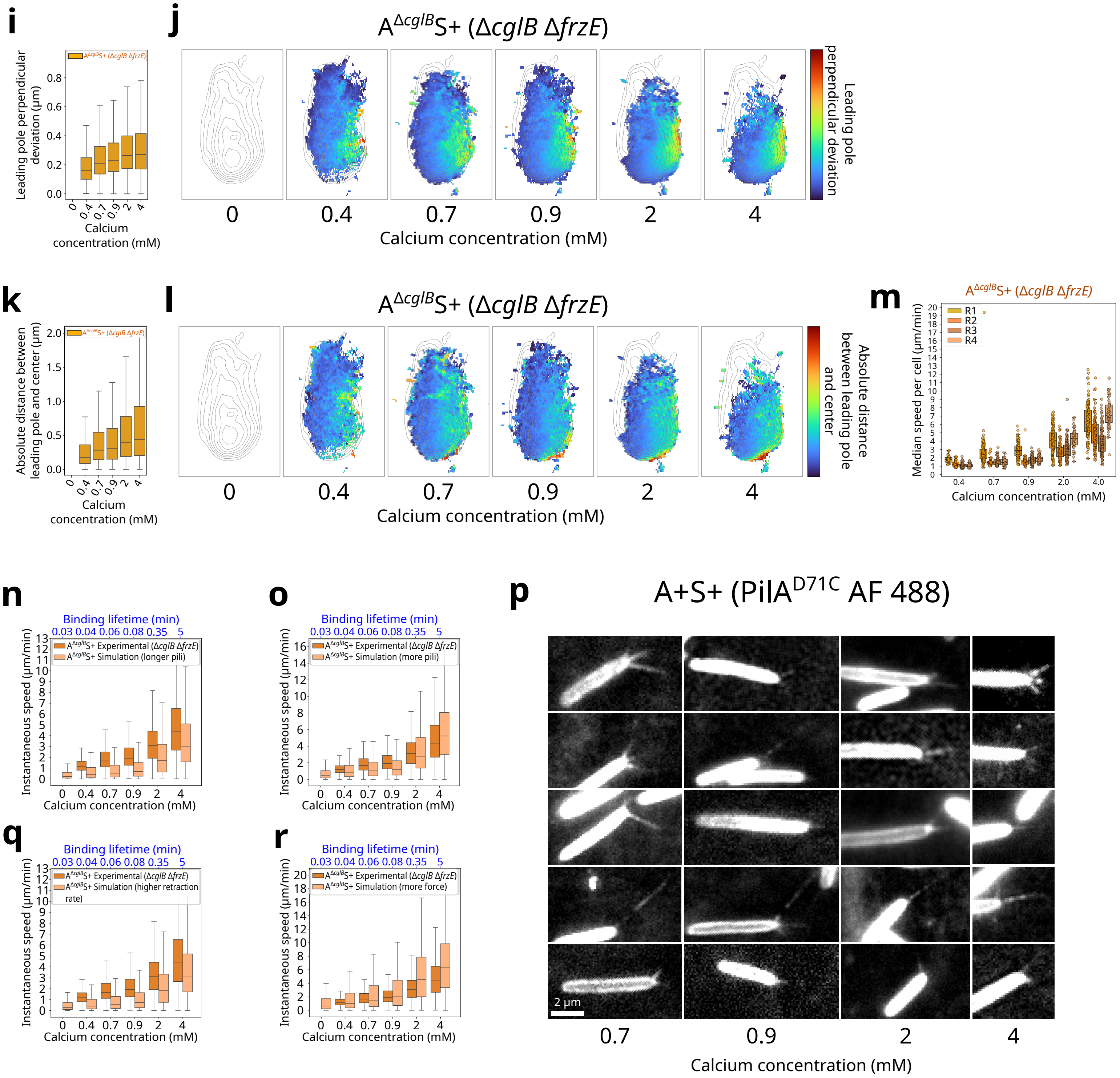

i
j
A^ΔcglB S+ (ΔcglB ΔfrzE)
Leading pole perpendicular deviation (µm)
Leading pole perpendicular deviation
0
0.4
0.7
0.9
2
4
Calcium concentration (mM)
k
l
A^ΔcglB S+ (ΔcglB ΔfrzE)
Absolute distance between leading pole and center (µm)
Absolute distance between leading pole and center
m
Median speed per cell (µm/min)
R1
R2
R3
R4
n
o
q
r
Binding lifetime (min)
0.03 0.04 0.06 0.08 0.35 5
Instantaneous speed (µm/min)
A^ΔcglB S+ Experimental (ΔcglB ΔfrzE)
A^ΔcglB S+ Simulation (longer pili)
A^ΔcglB S+ Simulation (more pili)
A^ΔcglB S+ Simulation (higher retraction rate)
A^ΔcglB S+ Simulation (more force)
p
A+S+ (PilA^D71C AF 488)
2 µm

#### Supplementary Figure 4

**(a)** Fluorescence images acquired in HILO mode (488 nm laser) on a chitosan-coated microfluidic device, showing PilA$^{D71C}$ AF488 labeled type IV pili in A$^{-}$S$^{+}$ (Δ*aglQ*) cells across calcium concentrations.

**(b-c)** Mean number of static AglZ-YFP clusters per cell across calcium concentration for experimental strains replicates. Error bars show ± SD within each replicate. (b) A$^{+}$S$^{-}$ (Δ*pilA*, red) replicates. R1 : n = 4, 15, 16, 19, 11, 12 cells at 0, 0.4, 0.7, 0.9, 2, 4 mM. R2 : n = 9 , 8, 9, 9, 5, 5 cells at 0, 0.4, 0.7, 0.9, 2, 4 mM. (c) A$^{+}$S$^{+}$ (green) replicates. R1 : n = 2, 63, 71, 82, 64, 39 cells at 0, 0.4, 0.7, 0.9, 2, 4 mM. R2 : n = 5, 45, 56, 41, 27, 14 cells at 0, 0.4, 0.7, 0.9, 2, 4 mM.

**(d)** Examples of automated kymographs of AglZ-YFP signal in A$^{+}$S$^{-}$ (Δ*pilA*) cells across calcium concentrations.

**(e)** Examples of automated kymographs of AglZ-YFP signal in A$^{+}$S$^{+}$ cells across calcium concentrations.

**(f-g)** Mean number of static AglZ-YFP clusters per cell according to cell speed for experimental strains replicates. Error bars show ± SD within each replicate. (f) A$^{+}$S$^{-}$ (Δ*pilA*, red) replicates. R1 : n = 77 cells. R2 : n = 45 cells. (g) A$^{+}$S$^{+}$ (green) replicates. R1 : n = 321 cells. R2 : n = 188 cells.

**(h)** Examples of automated kymographs of AglZ-YFP signal in A$^{\Delta cglB}$S$^{+}$ (Δ*cglB*) cells across calcium concentrations.

**(i)** Distribution of the leading pole perpendicular deviation for A$^{\Delta cglB}$S$^{+}$ (Δ*cglB* Δ*frzE*) cells across calcium concentrations. N = 4 replicates, n = – , – , 29, 67, 145, 190 cells at 0, 0.4, 0.7, 0.9, 2, 4 mM.

**(j)** UMAP projections of the full 48-dimensional behavior feature set for A$^{\Delta cglB}$S$^{+}$(Δ*cglB* Δ*frzE*) cells across calcium concentration. Rather than showing raw point density, each bin in the two-dimensional map is color-coded by the average normalized perpendicular deviation of the leading pole for all behavior segments falling into that bin. Iso-contours represent the combined behavioral density of all four strains pooled at the calcium concentration of 0.9 mM.

**(k)** Distribution of the absolute distance between leading pole and center for A$^{\Delta cglB}$S$^{+}$ (Δ*cglB* Δ*frzE*) cells across calcium concentrations. N = 4 replicates, n = – , 155, 252 , 236, 335, 372 cells at 0, 0.4, 0.7, 0.9, 2, 4 mM.

**(l)** UMAP projections of the full 48-dimensional behavior feature set for A$^{\Delta cglB}$S$^{+}$ (Δ*cglB* Δ*frzE*) cells across calcium concentration. Rather than showing raw point density, each bin in the two-dimensional map is color-coded by the absolute distance between leading pole and center for all behavior segments falling into that bin. Iso-contours overlay the combined occupancy (density) across all four strains and calcium concentrations.

**(m)** Median instantaneous speed per cell for $A^{\Delta cglB}S^{+}$ (Δ*cglB* Δ*frzE*) strains across calcium concentrations and replicates. R1 : n = –, 28, 99, 76, 99, 129 cells at 0, 0.4, 0.7, 0.9, 2, 4 mM. R2 : n = – , 35, 34, 33, 61, 96 cells at 0, 0.4, 0.7, 0.9, 2, 4 mM. R3 : n = –, 54, 43, 53, 64, 90 cells at 0, 0.4, 0.7, 0.9, 2, 4 mM. R4 : n = – , 27, 43, 39, 46, 51 cells at 0, 0.4, 0.7, 0.9, 2, 4 mM.

**(n)** Instantaneous speed per cell for experimental $A^{\Delta cglB}S^{+}$(Δ*cglB* Δ*frzE,* orange) and simulated $A^{\Delta cglB}S^{+}$ with longer pili (peach) across calcium concentrations or binding lifetime. Simulated $A^{\Delta cglB}S^{+}$ (Δ*cglB* Δ*frzE*) : n = 100 simulations per binding lifetime conditions. Sample size for experimental $A^{\Delta cglB}S^{+}$(Δ*cglB* Δ*frzE*) is the same as in Figure 4n.

**(o)** Instantaneous speed per cell for experimental $A^{\Delta cglB}S^{+}$ (Δ*cglB* Δ*frzE,* orange) and simulated with more pili (peach) across calcium concentrations or binding lifetime. Simulated $A^{\Delta cglB}S^{+}$ (Δ*cglB* Δ*frzE*) : n = 100 simulations per binding lifetime conditions. Sample size for experimental $A^{\Delta cglB}S^{+}$ (Δ*cglB* Δ*frzE*) is the same as in Figure 4n.

**(p)** Fluorescence images acquired in HILO mode (488 nm laser) on a chitosan-coated microfluidic device, showing $PilA^{D71C}$ AF488 labeled type IV pili in $A^{+}S^{+}$ cells across calcium concentrations.

**(q)** Instantaneous speed per cell for experimental $A^{\Delta cglB}S^{+}$ (Δ*cglB* Δ*frzE,* orange) and simulated $A^{\Delta cglB}S^{+}$ with higher retraction rate (peach) across calcium concentrations or binding lifetime. Simulated $A^{\Delta cglB}S^{+}$ (Δ*cglB* Δ*frzE*) : n = 100 simulations per binding lifetime conditions. Sample size for experimental $A^{\Delta cglB}S^{+}$ (Δ*cglB* Δ*frzE*) is the same as in Figure 4n.

**(r)** Instantaneous speed per cell for experimental $A^{\Delta cglB}S^{+}$ (Δ*cglB* Δ*frzE,* orange) and simulated $A^{\Delta cglB}S^{+}$ with higher force (peach) across calcium concentrations or binding lifetime. Simulated $A^{\Delta cglB}S^{+}$ (Δ*cglB* Δ*frzE*) : n = 100 simulations per binding lifetime conditions. Sample size for experimental $A^{\Delta cglB}S^{+}$ (Δ*cglB* Δ*frzE*) is the same as in Figure 4n.

#### Supplementary Movies

https://osf.io/pgnws/?view_only=8318cfd9386f4cac91e7d752be7a47cc

**(Movie S1)** AglZ-YFP clusters tracking and classification. Example of an $A^+S^-$ cell expressing AglZ-YFP, imaged by TIRF. Individual clusters were detected based on fluorescence intensity, tracked across successive frames, and classified as static (magenta) or mobile (green) according to their displacement. Images were acquired every 1 s, and the movie is displayed at 50 frames per second. Playback corresponds to ~50× real time.

**(Movie S2)** Fluorescence imaging of $A^+S^+$ cells expressing AglZ-YFP (green clusters) and *E. coli* cells expressing HU-mCherry (red) in the three regions shown in Fig. 1c of the predation assay: large swarms (A), small swarms (B), and isolated cells (C). Images were acquired every 5 s, and the movie is displayed at 10 frames per second. Playback corresponds to ~50× real time.

**(Movie S3)** HILO/TIRF time-lapse of type IV pili (left) and AglZ-YFP (right) in $A^+S^+$ cells (AglZ-YFP PilA$^{D71C}$ Atto 565) on 0.75% agar pads. The movie shows the entire field of view. Images were acquired every 2 s, and the movie is displayed at 25 frames per second. Playback corresponds to ~50× real time.

**(Movie S4)** HILO/TIRF time-lapse of type IV pili (right) and AglZ-YFP (left) in $A^+S^+$ cells (AglZ-YFP PilA$^{D71C}$ Atto 565) on 0.75% agar pads. The movie shows representative examples of individual cells (A, B, C). Images were acquired every 2 s, and the movie is displayed at 25 frames per second. Playback corresponds to ~50× real time.

**(Movie S5)** HILO time-lapse of an $A^-S^+$ mutant (Δ*aglQ* PilA$^{D71C}$ Atto 565) on 0.75% agar pads. Images were acquired every 2 s, and the movie is displayed at 25 frames per second. Playback corresponds to ~50× real time.

**(Movie S6)** HILO/TIRF time-lapse of type IV pili and AglZ-YFP dynamics in $A^+S^+$ cells (AglZ-YFP PilA$^{D71C}$ Atto 565) inside a chitosan-coated microfluidic chamber with 0.9 mM $CaCl_2$. Panels A-C show representative cells with AglZ-YFP on the left and pili on the right. Images were acquired every 5 s, and the movie is displayed at 10 frames per second. Playback corresponds to ~50× real time.

**(Movie S7)** HILO/TIRF and phase-contrast time-lapse of type IV pili and AglZ-YFP dynamics in $A^+S^+$ cells (AglZ-YFP, PilA$^{D71C}$ Atto 565) shown in Fig. 2a, inside a chitosan-coated microfluidic chamber with 0.9 mM $CaCl_2$. The left panel merges phase contrast and AglZ-YFP, while the right panel shows pili. Images were acquired every 5 s, and the movie is displayed at 10 frames per second. Playback corresponds to ~50× real time.

**(Movie S8)** TIRF and phase-contrast time-lapse of AglZ-YFP in $A^+S^+$ cells inside a chitosan-coated microfluidic chamber with 0.9 mM $CaCl_2$. Panel A shows AglZ-YFP, panel B shows phase contrast, and panel C displays tracked trajectories of the leading pole, lagging pole, and center for each cell, color-coded by time. Images were acquired every 4 s, and the movie is displayed at 12.5 frames per second. Playback corresponds to ~50× real time.

**(Movie S9)** Full-field trajectories showing leading pole, lagging pole, and center positions color-coded by time, as in Fig. 2f-h. Cells were imaged in a chitosan-coated microfluidic chamber with 0.9 mM $CaCl_2$. Panels: (A) $A^+S^-$ (Δ*pilA* Δ*frzE*), (B) $A^-S^+$ (Δ*aglQ* Δ*frzE*), and (C) $A^+S^+$ (Δ*frzE*). Cells were tracked every 1 s, and the movie is displayed at 50 frames per second. Playback corresponds to ~50× real time.

**(Movie S10)** Full-field trajectories showing leading pole, lagging pole, and center positions under increasing $CaCl_2$ concentrations for $A^+S^-$ (Δ*pilA* Δ*frzE*) cells, color-coded by time. Cells were imaged in a chitosan-coated microfluidic chamber, tracked every 1 s, and the movie is displayed at 50 frames per second. Playback corresponds to ~50× real time.

**(Movie S11)** Full-field trajectories showing leading pole, lagging pole, and center positions under increasing $CaCl_2$ concentrations for $A^-S^+$ (Δ*aglQ* Δ*frzE*) cells, color-coded by time. Cells were imaged in a chitosan-coated microfluidic chamber, tracked every 1 s, and the movie is displayed at 50 frames per second. Playback corresponds to ~50× real time.

**(Movie S12)** Full-field trajectories showing leading pole, lagging pole, and center positions under increasing $CaCl_2$ concentrations for $A^+S^+$ (Δ*frzE*) cells, color-coded by time. Cells were imaged in a chitosan-coated microfluidic chamber, tracked every 1 s, and the movie is displayed at 50 frames per second. Playback corresponds to ~50× real time.

**(Movie S13)** Examples of simulated trajectories of gliding ($A^+S^-$) mutants, showing the leading pole, lagging pole, and cell center color-coded by time. The panel combines individual simulations, each representing a cell modeled as six disks (grey) connected by springs, with a constant propulsive force applied along its length (red disks). Positions were sampled every 1 s, and the movie is displayed at 50 frames per second. Playback corresponds to ~50× real time.

**(Movie S14)** Examples of simulated trajectories of twitching ($A^-S^+$, Δ*aglQ* Δ*frzE*) mutants, showing the leading pole, lagging pole, and cell center color-coded by time. The panel combines individual simulations, each representing a cell as six disks (grey) connected by springs, with propulsion applied only to the leading disk via pili (green lines). Pili binding lifetimes were set to reflect $CaCl_2$ concentration (0 mM = 0.03 min; 0.4 mM = 0.04 min; 0.7 mM = 0.06 min; 0.9 mM = 0.08 min; 2 mM = 0.35 min; 4 mM = 5 min). Positions were sampled every 1 s, and the movie is displayed at 50 frames per second. Playback corresponds to ~50× real time.

**(Movie S15)** Examples of simulated trajectories of the additive model ($A^+S^+$), merging $A^+S^-$ and $A^-S^+$ motility simulations, showing the leading pole, lagging pole, and cell center color-coded by time. The panel combines individual simulations, each representing a cell as six disks (grey) connected by springs, with propulsion applied to the leading disk via pili (green lines) whose binding lifetime varies with $CaCl_2$ concentration (0 mM = 0.03 min; 0.4 mM = 0.04 min; 0.7 mM = 0.06 min; 0.9 mM = 0.08 min; 2 mM = 0.35 min; 4 mM = 5 min), and a constant propulsive force along its length (red disks). Positions were sampled every 1 s, and the movie is displayed at 50 frames per second. Playback corresponds to ~50× real time.

**(Movie S16)** TIRF time-lapse of AglZ-YFP in $A^+S^-$ (Δ*pilA)* cells inside a chitosan-coated microfluidic chamber under increasing $CaCl_2$ concentrations. Images were acquired every 4 s, and the movie is displayed at 12.5 frames per second. Playback corresponds to ~50× real time.

**(Movie S17)** TIRF time-lapse of AglZ-YFP in $A^+S^+$ cells inside a chitosan-coated microfluidic chamber under increasing $CaCl_2$ concentrations. Images were acquired every 4 s, and the movie is displayed at 12.5 frames per second. Playback corresponds to ~50× real time.

**(Movie S18)** Full-field trajectories showing leading pole, lagging pole, and center positions under increasing $CaCl_2$ concentrations for $A^{\Delta cglB}S^+$ (Δ*cglB* Δ*frzE*), color-coded by time. The ΔfrzE background was used to suppress cell reversals and facilitate quantitative trajectory analysis. Cells were imaged in a chitosan-coated microfluidic chamber, tracked every 1 s, and the movie is displayed at 50 frames per second. Playback corresponds to ~50× real time.

**(Movie S19)** TIRF time-lapse of AglZ-YFP in $A^{\Delta cglB}S^+$ (Δ*cglB*) cells inside a chitosan-coated microfluidic chamber under increasing $CaCl_2$ concentrations. This strain retains Frz-dependent reversals and was used to visualize dynamic A-motility complexes in the absence of substrate anchoring. Images were acquired every 4 s, and the movie is displayed at 12.5 frames per second. Playback corresponds to ~50× real time.

**(Movie S20)** Examples of simulated trajectories of twitching mutants with lower damping ($A^{\Delta cglB}S^+$, Δ*cglB* Δ*frzE*), showing the leading pole, lagging pole, and cell center color-coded by time. The panel combines individual simulations, each representing a cell as six disks (grey) connected by springs, with propulsion applied only to the leading disk via pili (green lines) whose binding lifetime varies with $CaCl_2$ concentration (0 mM = 0.03 min; 0.4 mM = 0.04 min; 0.7 mM = 0.06 min; 0.9 mM = 0.08 min; 2 mM = 0.35 min; 4 mM = 5 min), and with reduced damping. Positions were sampled every 1 s, and the movie is displayed at 50 frames per second. Playback corresponds to ~50× real time.

#### Supplementary Material

| Species | Strain | Phenotype | Genotype | Reference |
|---|---|---|---|---|
| *M. xanthus* | TM9 | A+S+ ; AglZ-YFP | DZ2 *aglZ-YFP* (allelic recombination) | Mignot laboratory collection |
| *M. xanthus* | TM478 | A+S- ; AglZ-YFP | DZ2 *pilA*::tet *aglZ-YFP* (allelic recombination) | Mignot laboratory collection |
| *M. xanthus* | TM/SR1 | A-S+ ; AglZ-YFP | DZ2 Δ*cglB aglZ-YFP* (allelic recombination) | Mignot/Nollmann laboratory collection |
| *M. xanthus* | AM8 | A-S+ ; $PilA^{D71C}$ (maleimide-labellable pili) | DZ2 Δ*aglQ* $att_{Mx8}$::pSWU30-$P_{pilA}$-$pilA^{D71C}$ | This study |
| *M. xanthus* | AM1 | A+S+ ; AglZ-YFP ; $PilA^{D71C}$ (maleimide-labellable pili) | DZ2 $att_{Mx8}$::pSWU30-$P_{pilA}$-$pilA^{D71C}$ AglZ-YFP (allelic recombination) | This study |
| *M. xanthus* | RM384 | A+S+ ; $PilA^{D71C}$ (maleimide-labellable pili) | DZ2 $att_{Mx8}$::pSWU19-$P_{pilA}$-$pilA^{D71C}$ | Mignot laboratory collection |
| *M. xanthus* | TM726 | A+S+ ; ΔFrzE | DZ2 Δ*frzE* | Mignot laboratory collection |

| | | | | |
|---|---|---|---|---|
| *M. xanthus* | TM1536 | A+S- ; ΔFrzE | DZ2 *pilA*::tet Δ*frzE* | Mignot laboratory collection |
| *M. xanthus* | TM831 | A-S+ ; ΔFrzE | DZ2 Δ*aglQ* Δ*frzE* | Mignot laboratory collection |
| *M. xanthus* | TM604 | A-S+ ; ΔFrzE | DZ2 Δ*cglB* *frzE*::tet | Mignot laboratory collection |
| *M. xanthus* | TM146 | A-S+ | DZ2 Δ*aglQ* | Mignot laboratory collection |
| *E. coli* | | WT ; HU-mCherry | MG1655 HU-mCherry | Espeli laboratory collection |

#### Supplementary Table 1

Bacterial strains used in this study.

| **Plasmid** | **Genotype** | **Reference** |
|---|---|---|
| pSWU30-$P_{pilA}$-$pilA^{D71C}$ | pSWU30 to express $P_{pilA}$-$pilA^{D71C}$ variant at Mx8 phage *attB* site | Mignot laboratory collection |

#### Supplementary Table 2

Plasmid used in this study.

#### Single-cell tracking

We used LapTrack tracking [1] to link segmented cell masks across frames via a custom cost function combining four terms:

1. **Overlap cost**: $1 - \frac{intersection\ area}{area\ of\ mask_{t+1}}$
2. **Distance cost**: squared Euclidean distance between mask centroids
3. **Collinearity cost**: $1-|\cos\theta|$, where $\theta$ is the angle between the observed displacement vector and the cell's orientation
4. **Area cost**: $\frac{|A_t - A_{t+1}|}{\frac{1}{2}(A_t + A_{t+1})}$

Each term was equally weighted (all weights = 1), and any candidate link whose total cost exceeded 0.9 was rejected. We disabled automatic gap closing and division detection (gap_closing_max_frame_count = 0, splitting_cost_cutoff = 0), producing an initial set of track segments. After LapTrack assignment, we relabeled each track with a unique ID and wrote out a relabeled TIFF stack for visualization.

##### Fragmented‑track reconnection

To reconnect segments broken by temporary segmentation loss, we considered all segment endpoints that reappeared within a 10-pixel radius and within 30 frames. For each candidate pairing, we computed a simple score: $S=w_{dist}.d+w_{time}.\Delta t$, where $d$ is the Euclidean distance between the end and start centroids, $\Delta t$ is the frame gap, and $w_{dist}$ and $w_{time}$ are user‑defined weights. Weights were used to favor spatial closeness over temporal proximity, or vice versa (both set to 1 by default). For each gap, we chose the single lowest-S pairing to merge segments into continuous tracks.

Finally, we discarded all tracks shorter than five frames, performed a manual validation in Napari, and applied basic morphological/kinetic filters to remove spurious trajectories before downstream analysis. Resulting trajectories were reassigned with a unique ID, and we relabeled a TIFF stack for cell poles and center detection.

##### Cell poles and center detection

After relabeling each mask with its track ID, we recovered pole and center coordinates directly from the cell outline in every frame. First, the binary mask was skeletonized and cleaned to yield a single‑branch medial axis, which was then fit with a smooth spline to trace the full, possibly curved, length of the cell. Poles were initially estimated in two ways: by taking the spline endpoints extrapolated to the cell contour, and by locating minima in the contour curvature, and then reconciled by pairing each curvature-derived point with its nearest spline endpoint and averaging their coordinates when both were available (falling back to whichever estimate was present if one method failed). Finally, the cell's geometric center was computed as the midpoint along the spline's arc length. These subpixel-refined pole1, pole2, and center positions were appended to our centroid tracks for all downstream analyses.

**Cell poles and center preprocessing**

Before calculating any polarity or center features, we applied a standard two-step preprocessing pipeline to all pole and center trajectories. First, each raw (x,y,time) track was spatially resampled to a constant step length (0.025 µm by default), and its original timestamps were then interpolated onto the new points via dynamic-time-warping alignment to preserve the true temporal sequence. Second, to filter high-frequency noise from segmentation jitter and localization error, the rediscretized trajectories were smoothed with a Savitzky-Golay filter (with a window length of 11 points). These uniformly spaced, denoised tracks of the leading pole, lagging pole, and cell center then served as the input for all subsequent feature computations.

**Signed-displacement calculation**

Signed-displacement calculation then quantified how each cell moved relative to its poles. We first computed frame-to-frame center deltas (Δx, Δy) and, for each frame, the unit vector pointing from pole 1 to pole 2. By projecting the displacement vector onto this pole vector, we obtained a scalar "displacement amplitude" (the magnitude of motion along the cell's long axis) and a binary "sign" (+1 or -1) indicated whether pole 2 or pole 1 lay ahead in the direction of movement. Using these +1/-1 flags, we assigned each pole-specific measurement to the leading or lagging pole at each time point.

The resulting signs were smoothed with a 20-frame sliding-window mode filter to remove spurious flips that falsely suggest the poles have moved backwards.

**Inspection and filtering of trajectories**

Cell masks and their corresponding tracks were overlaid on the original images in Napari for visual inspection. We manually inspected each cell to confirm that its mask matched the underlying cell morphology and that its trajectory showed no tracking or segmentation artifacts. Only those cells passing this quality check were retained, and all others were excluded by filtering their track IDs out of the analysis DataFrame.

After manual filtering in Napari, we filtered out any trajectories that did not meet a set of morphological and kinetic criteria to eliminate potential artifacts. For each cell track as a whole, its true length had to remain stable (standard deviation less than twice the median), with a median length between 15 and 100 pixels, and its median center speed between 0 and 20 000 pixels/s. Then, at every time point, the instantaneous displacement had to fall within 0-500 pixels, the cell's aspect ratio (perimeter divided by length) had to stay between 2 and 2.4, and the perimeter deviation from its median value had to be under ten times its median standard deviation. Finally, instantaneous speeds at the cell center and at both poles were required to lie between 0 and 1 000 pixels/s. Only the data points that passed all of these filters were carried forward into the final analysis.

#### Features computation

For each smoothed and rediscretized trajectory (leading pole, lagging pole, or center), we computed the following windowed features, where window sizes refer to the number of rediscretized trajectory points (not time in seconds):

- **Angular dispersion**

  Computed in a 2 000-point sliding window centered on each timepoint. For every pair of consecutive trajectory points, we calculate the motion angle $\theta=\mathrm{atan2}(\Delta y,\Delta x)$. We then map each $\theta$ to a unit-circle vector $(\cos\theta,\sin\theta)$ and take the mean of those vectors. If all steps point the same way, the mean-vector's length R is close to 1; if headings are scattered, R is near 0. We report dispersion as $\mathrm{AngularDispersion}=1-R$, so 0 means straight motion, and values up to 1 indicate highly variable turning.

- **Auto-collinearity**
  Auto-collinearity was computed over 1000-point sliding window by comparing two displacement vectors: one from the window's start to its midpoint, and the other from the midpoint to its end. At each step, we calculated the cosine of the angle between these consecutive vectors, values near 1 indicate that the path segments are nearly collinear (straight), while lower or negative values reflect sharp bends or backtracking.

- **Directional autocorrelation**

  Directional autocorrelation was computed within a 2 000-point sliding window by first normalizing each displacement vector between consecutive points to unit length, averaging those unit vectors to obtain the window's mean-direction vector, and then taking the dot product of each individual unit vector with that mean. This descriptor quantified motion persistence in the cell's predominant direction of motion. Values near +1 indicate that steps closely follow the average direction (high persistence), whereas lower or negative values reflect frequent directional changes.

- **Directional bias index (DBI)**

  Directional bias index was defined over a forward-looking window of 500 points starting at each frame, and then assigned to the window's center. It quantified the ratio of net displacement (straight-line distance from the first to the last point of the window) to total path length (sum of all stepwise distances). Values near 1 indicate almost all motion is directed along one axis, whereas values approaching 0 denote meandering or back-and-forth movement.

- **Displacement-vector crossing rate**

  Within each window (500 points), we treat every consecutive-point displacement as a 2D vector and determine whether it lies "left" or "right" of the window's net displacement (using the sign of the 2D cross-product). We then count how many times the left/right sign flips over the window and divide by the window length. Higher values indicate more frequent oscillations or weaving in the trajectory.

- **Distances between coordinate pairs**

  At each time point, we compute the Euclidean distances between every pair of reference points—leading pole, lagging pole, and cell center—using their (x,y) coordinates. To place all trajectories into a common reference frame at each time point, we first align them via Principal Component Analysis (PCA) (so that their principal axes coincide) before measuring distances. The resulting values capture instantaneous spatial separations within the cell (e.g., how far the poles lie from the center or from each other).

- **Drift-to-diffusion ratio**

  Computed over each 2000 points sliding window as the ratio of net straight-line displacement to the mean squared deviation from the starting point (normalized by twice the window length). This unitless metric compares the magnitude of overall displacement to the typical spread of the path: higher values indicate that the cell's movement closely follows a straight chord, while lower values reflect more diffusive or exploratory wandering.

- **Entropy of motion**

  Entropy of motion was computed within a 2 000-point sliding window by first measuring the change in direction between one displacement vector and the next. Those angles were grouped into a fixed number of bins to form a simple histogram of how often each angle occurred. Shannon entropy of that distribution then reports how evenly spread the turning angles are: high entropy means turns are unpredictable and varied; low entropy means the cell sticks to just one or two preferred turning angles.

- **Local curvature**

  For each 2000 points sliding window, we compute the angle between each pair of successive displacement vectors (i.e., the instantaneous turning angle) and then take the average of those angles. This mean turning angle summarizes the path's "bendiness": values near 0° correspond to almost straight segments, whereas larger mean angles indicate sharper or more frequent curves.

- **Local radius of gyration**

  Within each 2000 points sliding window, we find the centroid of all trajectory points, calculate each point's squared distance to that centroid, and take the square root of their mean. This quantifies how widely the path fans out around its local center: higher values reflect more dispersed, exploratory motion.

- **Perpendicular deviations**

  Perpendicular deviations were computed by defining, for each 2 000-point window, the straight-line segment joining the window's first and last trajectory coordinates. For every trajectory point between those endpoints, we computed its perpendicular distance to this line and then averaged these distances to produce a single "perpendicular deviation" value per window. Because this metric quantified how far the cell deviated laterally from its overall direction of travel, larger values indicated more side-to-side wandering, whereas values near zero indicated movement closely following a straight path.

- **Mean projected displacement**

Mean projected displacement was computed within a 1 000-point sliding window by first averaging all unit displacement vectors in the window to obtain a local mean-direction vector, then projecting each instantaneous displacement vector onto that mean (via their dot product). This yielded a signed scalar at each step: positive values indicated net forward progress along the cell's prevailing direction, whereas values near zero corresponded to primarily lateral motion.

- **Segmented directional correlation**

  Segmented directional correlation was computed within a 2 000-point sliding window by first summing all displacement vectors in the window's first half and separately summing those in its second half to obtain two net-displacement vectors. We then normalized each to unit length and took their dot product (cosine similarity). Values near 1 indicate that the cell's overall direction in the first half closely matched that in the second half (high coherence), whereas lower values signify a substantial shift in trajectory between the two segments.

- **Straightness index**

  Computed identically to DBI but over a centered window (half before, half after the current point), this ratio of net displacement to total path length provides a validation check: it should closely match the forward-looking DBI when the windows are symmetric.

- **Tortuosity**

  Tortuosity was defined as the ratio of the total path length traveled (the sum of distances between each pair of consecutive points over a 2000 points sliding window) to the straight-line distance between the window's start and end points so that values above 1 reflected increasingly winding trajectories.

- **Trajectory angles**
  Trajectory angles were computed by measuring the signed angle between displacement vectors from the start to the midpoint and from the midpoint to the end of each 2000 points sliding window, with positive and negative values indicating leftward or rightward turns, respectively.

#### Feature preprocessing and UMAP embedding

Feature preprocessing and UMAP embedding were performed as follows. First, we assembled a unified feature matrix by selecting our curated set of morphodynamic descriptors and metadata, then applied a series of deterministic, feature-specific transformations in Python (log-scaling of distance and ratio metrics, absolute-value correction of signed angles, and a mild nonlinear “sharpening” of collinearity scores) to bring each descriptor into a comparable numeric range. We then removed any rows containing NaN or infinite values. Next, we standardized the resulting matrix using a power transform (to symmetrize distributions and reduce skew) and used the UMAP algorithm (umap-learn) with n_neighbors=50, min_dist=0.9, metric="euclidean", init="spectral", and n_components=2 to compute a two-dimensional embedding.

#### AglZ-YFP detection and tracking

Focal adhesion complexes' fluorescence spots were detected and tracked in three main steps. First, we applied the Spotiflow deep-learning detector [2] to each frame of the drift-corrected fluorescence stack, retaining only spot calls with probability ≥0.3 and recording their subpixel (x,y) positions and frame indices. Next, we linked these detections into trajectories using TrackPy’s algorithm [3]: spots were allowed to jump up to 2 px between frames, could disappear for up to 10 frames, and reappear within that radius, and any track shorter than 5 frames was discarded. Finally, to prepare for downstream feature extraction, each spot trajectory was temporally interpolated to fill small gaps, then smoothed with a Savitzky-Golay filter (window = 11 frames) to filter localization noise before computing motility descriptors

- **AglZ-YFP clusters classification**

After smoothing, we applied the same trajectory rediscretization and computed some of the features described above for cell trajectories (tortuosity, entropy of motion, local curvature, crossing rate, perpendicular deviation, and persistence score), adding instantaneous velocity to the list. We standardized each feature to zero mean and unit variance, then fit a two-component Gaussian Mixture Model (scikit-learn) to separate “static” from “mobile” clusters based on their combined motility signatures.

- **AglZ-YFP clusters reassignment to cell masks**

We then linked each detected spot back to its corresponding cell by finding the nearest cell mask at each frame and assigning that cell's track ID to the spot. Specifically, for every spot's (x,y) coordinate and time point, we queried the masks for that frame within a fixed search radius and collected the IDs of any candidate masks. We then computed the Euclidean distance from the spot to each candidate mask's pixels and selected the mask whose centroid lay closest. By doing this for every spot and retaining only a single spot-cell pairing, we produced the final DataFrame in which each spot event carries its assigned cell's track ID. Finally, to ensure that we only analyzed adhesion events along the basal cell body, we computed the Euclidean distance between each spot and the corresponding cell's leading-pole coordinate. We then discarded any spot whose detection lay less than 5 px from the leading pole.

#### Simulations

All simulation parameters were drawn from published values when available and are summarized in Supplementary Table 3.

- **Bacterium architecture and dynamics**

The bacterium was represented as a linear chain of $N = 6$ circular disks of radius 0.5 µm [4], connected via linear springs of stiffness $k = 3{\times}10^3$ pN/µm to maintain structural cohesion [5]. To preserve shape, we introduced a bending penalty between every triplet of adjacent disks, applying corrective forces based on angular deviations from a straight configuration. This bending stiffness was controlled by a coefficient of $10^3$ pN [5].

The system evolved over time using velocity Verlet integration with a time step $\Delta t = 1/60$ min (i.e., 1 s), over 500 simulation steps unless stated otherwise. Damping forces proportional to velocity ($\gamma = 8$ pN·min/µm) represented surface friction and viscous resistance from the surrounding environment. This value lies within a factor of 3-4 of the 30 pN·min/µm inferred by Harvey et al., which is reasonable given the indirect nature of both estimates [6].

- **Pilus-based propulsion**

Pili were modeled as seven dynamic tethers extending from the leading disk, each with a rest length of ℓrest = 5 µm and a spring stiffness of kpili = 5000 pN/µm [7–9]. In each binding event, the actual pilus-substrate distance was sampled uniformly between 0.33 x ℓrest and 3 x ℓrest. Each pilus could bind stochastically to a surface site located within a forward-facing angular cone defined by a central direction (forward_angle = 0 rad) and a half-angle

(angle_of_view = π/1.5). Binding attempts occurred with a fixed probability per unit time, controlled by the binding_rate parameter (default: 1 event per minute) [10]. Upon successful binding, the pilus established a tether between the leading disk and the binding site. As the pilus retracts at the specified retraction rate (30 μm/min, [11]), its instantaneous length becomes shorter than this rest length and it generates a linear spring pulling force $F = k_{pili} \times (\ell_{rest} - \ell_{current})$. This pulling force was capped at a maximum value (max_motor_force, default: 70-200 pN depending on the simulated genetic background) to mimic motor stalling [12]. A pilus unbound under three conditions: (1) when its current length reaches zero (or a small numerical tolerance), reflecting complete retraction of the pilus; (2) when its binding duration exceeded a randomly sampled lifetime centered around the parameter binding_lifetime (default: 5 min) [10], or (3) if the binding site exited the allowed angular view due to changes in cell orientation or movement. All three events were evaluated at each time step to determine unbinding. At every simulation step, the net pilus force was computed by summing the contributions of all currently bound pili acting on the leading disk.

- **FACS-based adhesion**

Focal adhesion complexes (FACS) were modeled as adhesive contact points that were deposited by the leading disk when it traveled beyond a threshold distance from the most recent adhesion site. This distance threshold was drawn from a log-normal distribution centered on a nominal value (typically 2 μm), introducing stochasticity in adhesion spacing. Once deposited, each FACS was static and acted on any disk that came within interaction range (≤ 2 × disk radius). It exerted two distinct forces on each overlapping disk. First, a restoring spring force was directed from the disk toward the FACS point; its magnitude scaled with the distance between the disk and the adhesion site, with stiffness set by the parameter k_facs (default: 20 pN/μm). Second, a directional pushing force was applied along the local axis of the cell at the disk's position. This axis was estimated using the vector between neighboring disks (from disk *i-1* to *i+1*, or an appropriate approximation at the ends of the chain), and was scaled by a constant amplitude FACS_force (default: 20 pN per FACS, totalling ~60 pN traction per cell [13]). Together, these forces allowed FACS to anchor the cell to the substrate while biasing its movement in the direction of the chain, mimicking directional traction mechanisms observed in A-motility.

- **Simulation outputs**

At each time step, we recorded the positions of all disks (trajectory), pilus states (e.g., bound/unbound, binding time, force), and FACS point coordinates. These trajectories were later used to extract instantaneous speeds and morphodynamic descriptors for behavioral analysis as described for experimental data.

- **Calcium-dependent parameter sweeps**

To capture the graded effect of extracellular $CaCl_2$ on pilus behavior, we first assigned each tested pilus parameter (e.g. binding lifetime, retraction rate, stall force) a minimum value at 0 mM and a maximum value at 4 mM. Then, for each intermediate $CaCl_2$ level (0.4, 0.7, 0.9, 2 mM), we normalized the concentration onto a 0-1 scale and exponentially spaced the parameter between its minimum and maximum according to that normalized range.

- **Mutant-specific parameter variations**

To simulate distinct genetic backgrounds or calcium-dependent behaviors, we systematically varied a subset of biophysical parameters. The table below summarizes the key parameters and the conditions under which they were modified:

| **Category** | **Parameter** | **Description** | **Default value** | **Adjusted Values** | **Literature values** |
|---|---|---|---|---|---|
| Core simulation parameters | N | Number of disks per bacterium | 6 | Constant | N/A |
| | radius | Radius of each disk (μm) | 0.5 | Constant | ~0.4 (Pelling et al. 2005) |
| | dt | Time step (min) | 1 / 60 | Constant | N/A |
| | time_steps | Total number of simulation steps | 500 | Constant | N/A |
| | k | Spring stiffness between | $3 \times 10^3$ | Constant | $10^4$ (Janulevicius et al. 2010) |

| | | | | | |
|---|---|---|---|---|---|
| | | disks (pN/μm) | | | |
| | stiffness_coefficient | Bending stiffness coefficient (pN·μm/radian) | $10^3$ | Constant | 1 - $10^3$ (Janulevicius et al. 2010) |
| | damping_coefficient | Viscous drag coefficient (pN·min/μm) | 8 | 2.5 for $A^-S^+$ (ΔcglB ΔfrzE) with less damping simulations | 30 (Harvey et al. 2011) |
| Pili parameters | rest_length | Initial pilus length at binding (μm) | 5 | 10 for $A^-S^+$ (ΔcglB ΔfrzE) with longer pili | 2-6 (Treuner-Lange et al. 2020) |
| | binding_rate | Pilus binding rate (event/min) | 1 | Constant | 1 (Koch et al. 2021) |
| | angle_of_view | Pilus angular binding range (rad) | π / 1.5 | Constant | This work |
| | forward_angle | Direction of pilus projection (rad) | 0 | Constant | N/A |
| | max_pili_count | Maximum number of pili per cell | 7 | 14 for $A^-S^+$ (ΔcglB ΔfrzE) with more pili | 6.5±3.0 (Bulyha et al. 2009) |
| | | | | 0 for $A^+S^-$ | |

| | | | | | |
|---|---|---|---|---|---|
| | binding_lifetime | Maximum pilus binding time (min) | 5 | 0.03-5 to mimic calcium dependence in $A^-S^+$ and A+S+ | 0.15 (Koch et al. 2021) |
| | max_motor_force | Maximum pulling force per pilus (pN) | 70 | 200 for $A^-S^+$ (ΔcglB ΔfrzE) with more force | 70-150 (Clausen et al. 2009) |
| | retraction_rate | Pilus retraction speed (µm/min) | 30 | 60 for $A^-S^+$ (ΔcglB ΔfrzE) with higher retraction rate | 30 (Skerker and Berg 2001) |
| | k_pili | Pilus spring constant (pN/µm) | $5 \times 10^3$ | Constant | $5 \times 10^3$ (Treuner-Lange et al. 2024) |
| FACS parameters | FACS_force | Force exerted by FACS adhesion (pN) | 20 | 0 for $A^-S^+$ | 12 (Balagam et al. 2014) |
| | k_facs | Spring constant of FACS pulling (pN/µm) | 20 | 0 for $A^-S^+$ | Empirically tuned |
| | FACS_threshold_distance | Distance to trigger new FACS site (µm) | 2 | Variable via log-normal sampling | This work |

##### Supplementary Table 3 : Default parameters for simulations

- **Parameter variability**

To mimic cell-to-cell heterogeneity, a subset of biophysical parameters was independently perturbed for each simulated trajectory using multiplicative log-normal noise ($\sigma$ = 50%). These included pilus-related parameters (e.g., rest_length, binding_rate, retraction_rate, max_pili_count, max_motor_force), as well as FACS-specific parameters (e.g., FACS_force, k_facs, and FACS_threshold_distance). The discrete parameter corresponding to the number of pili (max_pili_count) was rounded to the nearest integer after sampling.
